# Cell type-Specific *In Vivo* Proteomes with a Multi-copy Mutant Methionyl t-RNA Synthetase Mouse Line

**DOI:** 10.1101/2024.11.28.625838

**Authors:** Rodrigo Alvarez-Pardo, Susanne tom Dieck, Kristina Desch, Belquis Nassim Assir, Cristina Olmedo Salinas, Riya S. Sivakumar, Julian D. Langer, Beatriz Alvarez-Castelao, Erin M. Schuman

## Abstract

The functional diversity of cells is driven by the different proteins they express. While improvements in protein labeling techniques have allowed for the measurement of proteomes with increased sensitivity, measuring cell type-specific proteomes *in vivo* remains challenging. One of the most useful pipelines is bioorthogonal non-canonical amino acid tagging (BONCAT) with the MetRS* system, consisting of a transgenic mouse line expressing a mutant methionyl-tRNA synthetase (MetRS*) controlled by Cre recombinase expression. This system allows for cell type-specific labeling of proteins with a non-canonical amino acid (azidonorleucine, ANL), which can be subsequently conjugated to affinity or fluorescent tags using click chemistry. Click-modified proteins can then be visualized, purified and identified. The reduction in sample complexity allows for the detection of small changes in protein composition. Here we describe a multicopy MetRS* mouse line (3xMetRS* mouse line), which exhibits markedly enhanced ANL protein labeling, boosting the sensitivity and temporal resolution of the system and eliminating the need for working under methionine depletion conditions. Cell type-specific *in vivo* labeling is possible even in heterozygous animals, thus offering an enormous advantage for crossing the line into mutation and disease-specific backgrounds. Using the 3xMetRS* line we identified the *in vivo* proteome of a sparse cell population - the dopaminergic neurons of the olfactory bulb and furthermore determined newly synthesized proteins after short labeling durations following a single intraperitoneal ANL injection.

## Introduction

Cellular protein composition and how it is modified in response to physiological and pathological signals and events is key to understanding how cells function. It is clear that the same extra-or intracellular signals can lead to distinct proteomic responses in different cell types (Borisova *et al*, 2024; Hrvatin *et al*, 2018). Indeed, bulk analyses of proteomes can “average out” intrinsic cellular differences as well as cell type-specific differences in responses. One way to obtain cell type-specific proteomes exploits cell type-specific markers and fluorescent labeling, followed by enrichment using fluorescence-activated cell sorting (FACS) for subsequent MS studies. A drawback of this approach, however, is the loss of processes and compartments, such as dendrites or axons, during sample processing (Binek *et al*, 2019; Wiegleb *et al*, 2022). In contrast, BioID/TurboID-based methods are explicitly designed to identify proteins from targeted subcellular regions. Since protein populations are labeled in a proximity-dependent manner at a specific time point this method includes spatial and temporal information but does not distinguish between newly synthesized vs. pre-existing proteins (May *et al*, 2020).

Bio-orthogonal methods based on the use of artificial amino acid analogs as baits for protein visualization and purification are particularly amenable to the study of cell type-specific proteomes (Dieterich *et al*, 2006; Kiick *et al*, 2002). The incorporation of artificial amino acids into proteins can be genetically controlled to enable cell type-specific labeling (Alvarez-Castelao *et al*, 2017; Evans *et al*, 2024). For instance, the expression of a mutant methionyl-tRNA synthetase with an enlarged methionine recognition pocket (L274G point mutation; hereafter referred to as MetRS*) allows the charging of the methionine analog azidonorleucine (ANL) to the corresponding methionine tRNA. ANL has a slightly larger structure compared to methionine. This size difference is sufficient to exclude ANL recognition by the endogenous MetRS; thus, ANL is not incorporated into proteins in cells where MetRS* is not present (Tanrikulu *et al*, 2009). When MetRS* expression is driven by cell type-specific promoters, it enables specific labeling of proteomes with ANL and, after tissue lysis or fixation, derivatization of the azide group present in ANL with an alkyne by click chemistry. The alkyne can be used to visualize cell type-specific proteomes by immunofluorescence (FUNCAT) or Western blot (BONCAT) and to purify these proteomes. The MetRS* system overcomes the above mentioned challenges: it is possible to obtain cell type-specific proteomes without losing proteins located in dendrites or axons (Perez *et al*, 2021b). Additionally, the time window of protein synthesis is determined by the exposure time of ANL, and protein degradation or subcellular location of ANL-labeled proteins is not altered, using this method, it is even possible to measure protein half-lives (Alvarez-Castelao *et al*., 2017; tom Dieck *et al*, 2015).

Based on this system, we originally created a mouse model carrying a knock-in of the MetRS* gene fused to GFP by a P2A peptide under an enhanced actin promoter. Expression of the cassette is switched on by Cre-dependent excision of a transcriptional stop. Crossing this mouse line with cell type-specific Cre drivers allows for cell type-specific protein labeling and its subsequent visualization, purification, and identification. Given the considerable number of Cre drivers, the described system can be used to study cell type-specific proteins from almost any tissue in any field. Proteins can be labeled with ANL in live animals or *in vitro*. Using this first generation of mice, we demonstrated that it is possible to purify and identify cell type-specific proteomes allowing for the identification of the respective cell type of origin. Furthermore, we identified proteins whose expression in the excitatory hippocampal neurons is modified in response to an enriched environment, a well-established paradigm for synaptic plasticity enhancement (Alvarez-Castelao *et al*., 2017). It is also possible to identify changes in protein expression in models of prion diseases (Albert-Gasco *et al*, 2024). However, we observed limitations when we aimed to obtain proteomes from relatively low abundance cell types. Similarly, the identification of proteins labeled for short periods was not possible in brain tissue using this mouse model. Furthermore, given the line expresses a single copy of the MetRS*, we likely observed substantial competition with methionine incorporation by the endogenous MetRS. Indeed, to obtain good *in vivo* labeling, a low methionine diet and homozygous MetRS* alleles were required. The need for homozygosity complicates the crossing of this mouse line into a disease or any other background of interest. To overcome these limitations, we describe here a second generation of the MetRS* mouse tool designed to increase the expression of the mutant enzyme. The new design of the line boosts ANL incorporation into proteins, allowing for the isolation and identification of low-number neuronal populations, such as the dopaminergic neurons, and proteins synthesized in the excitatory neurons of the cortex after just 3 hours of a single intraperitoneal-IP-injection of ANL.

## Results

### The 3xMetRS* mouse line

In order to achieve more efficient ANL labeling to purify proteomes from sparse cell populations or from proteomes after short ANL labeling times, we focussed on improving the expression of MetRS*. We therefore modified and optimized the allele design of the previous MetRS* mouse line (termed 1xMetRS* here). We maintained the overall strategy of a Cre-dependent excision of a floxed transcriptional stop to switch on transcription of a cassette with MetRS* and GFP in the ROSA26 locus (Fig. 1A). Whereas the 1xMetRS* line carried a GFP sequence fused to the MetRS* coding region, separated by the self-cleaving 2A peptide sequence (P2A) (Fig. 1B), the new MetRS* mouse line described here (hereafter referred to as 3xMetRS*) expresses a cassette carrying three copies of MetRS* coding sequences and one copy of GFP per allele, with all four parts separated by sequences coding for different self-cleaving 2A peptides (Fig. 1C). As this expression cassette codes for a very large mRNA (and protein before separation of the single units by self-cleavage), we added an mRNA stabilization element (WPRE) to boost expression of the exogenous mRNA. In addition, we shifted the GFP to the C-terminus of the encoded protein (Fig. 1C). As such, we ensured that if GFP is detected, MetRS* is also translated. Mice carrying the 3xMetRS* allele and the Cre::3xMetRS* lines used in this work were viable and fertile, and both heterozygous and homozygous mice showed no obvious behavioral differences compared to wild type mice. We use the MetRS* copy numbers to refer to the genotype: 1xMetRS* and 2xMetRS* when studying heterozygous or homozygous animals, respectively, in the previously described mouse line, and 3xMetRS* or 6xMetRS* when using heterozygous or homozygous animals of the new line (Fig. 1).

**Figure 1.**
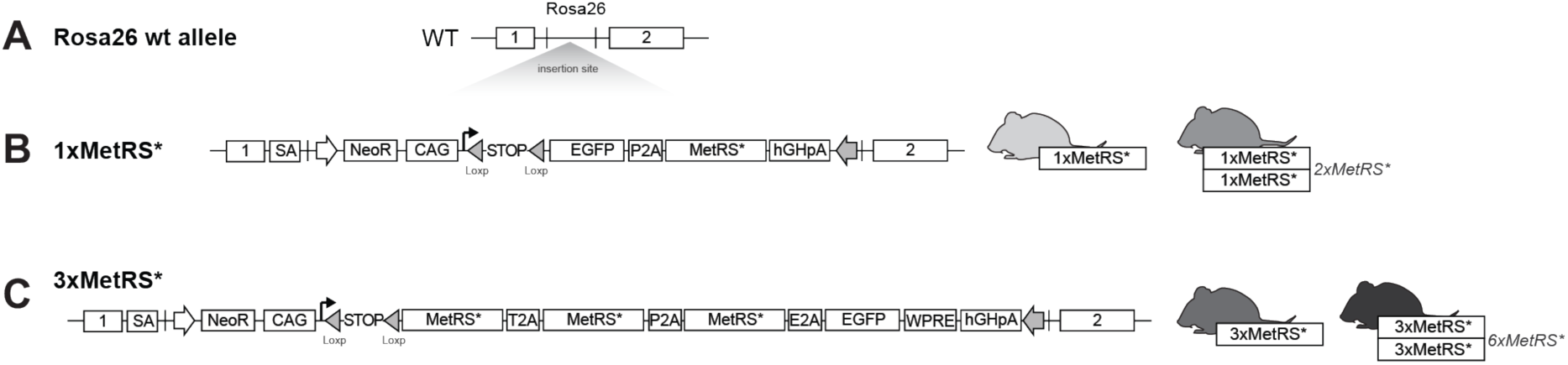
Comparison of the two mouse lines for ANL cell type-specific protein labeling. (**A**) Insertion of the MetRS* cassette in the ROSA26 locus. (**B**) Scheme of the cassette introduced in the first generation of the floxed STOP MetRS* mouse line. The right part shows mice expressing one (1xMetRS*, heterozygous animals) or two (2xMetRS*, homozygous animals) copies of the transgene after recombination by Cre. (**C**) Scheme of the cassette introduced in the second generation floxed MetRS*, showing all the introduced genes separated by 2A peptides, and the mRNA stabilization element (WPRE) at the end of the cassette. The right part shows mice expressing three (3xMetRS*, heterozygous animals) or six (6xMetRS*, homozygous animals) copies of the gene after recombination by Cre.

### Neuronal ANL labeling *in vitro* with different MetRS* copy numbers

To analyze the utility of increasing the MetRS* copy number, we compared ANL incorporation between the original and second generation lines. First, we evaluated whether there was increased ANL labeling in the 3xMetRS* line compared to the 1xMetRS* line *in vitro*. We crossed animals carrying the respective alleles to the Nex-Cre driver line, driving Cre expression to excitatory neurons (Goebbels *et al*, 2006), and performed experiments on cultured cortical neurons derived from the offspring. Experiments in culture have the advantage that ANL labeling is direct and circumvents any variability arising from ANL administration or uptake into the brain. Comparison of 2-hour ANL labeling in either 1xMetRS* or 3xMetRS* in excitatory neurons (Nex-Cre::1xMetRS* and Nex-Cre::3xMetRS*) evaluated by FUNCAT showed that ANL labeling was approximately three times higher in neurons expressing 3xMetRS* (Fig. 2A, B) (tom Dieck *et al*., 2015). FUNCAT experiments in the presence of the protein synthesis inhibitor anisomycin (Fig. 2C) confirmed that the observed fluorescent signal was due to ANL incorporation into newly synthesized proteins and not due to detection of free ANL or ANL-charged tRNAs. Interestingly, we observed that not only was ANL labeling increased in Nex-Cre::3xMetRS*, but also GFP expression was higher in the Nex-Cre::3xMetRS* line compared to Nex-Cre::1xMetRS*, despite the fact that there is one copy of GFP in both lines (Fig. 2D). Similarly, Western blot comparisons of ANL labeling in 1xMetRS* or 3xMetRS* cultured neurons by BONCAT also showed an increase in ANL labeling (Fig. 2E, F). To examine whether the observed increase in labeling with the 3xMetRS* line was also evident in tissue, we prepared acute slices from brain tissue of adult Nex-Cre::1xMetRS* or 3xMetRS* mice and performed *in vitro* slice metabolic ANL (2 h) labeling experiments. As was observed in the cultured neurons, FUNCAT labeling was significantly elevated in slices expressing the 3xMetRS* transgene (Fig. 2G).

**Figure 2.**
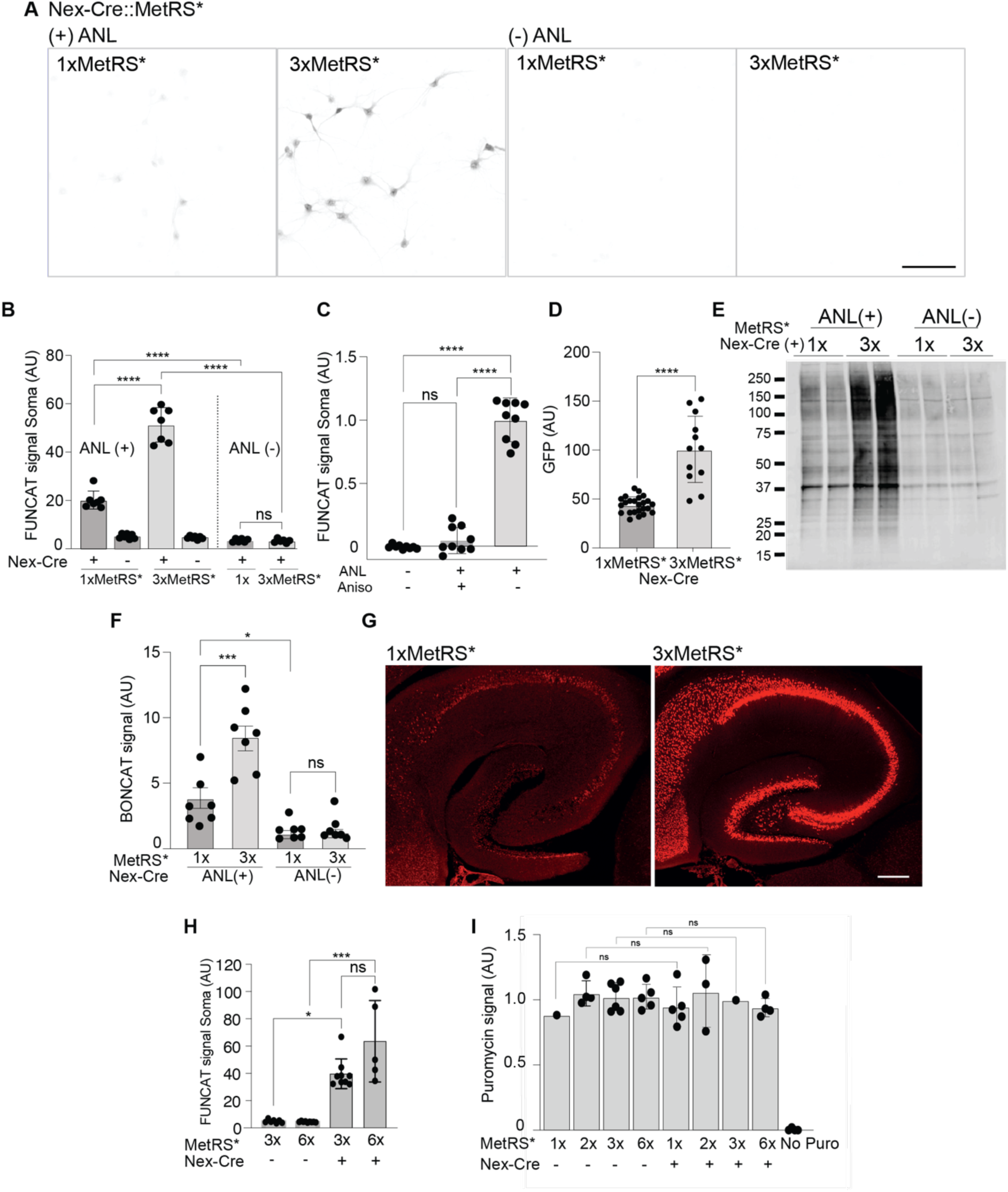
MetRS* copy number influence in ANL labeling. (**A**) Representative confocal images showing ANL labeling by FUNCAT after 2 h of incubation, in primary cortical neurons from Nex-Cre:1xMetRS* or Nex-Cre:3xMetRS* mice with (+) or without (-) ANL, showing an increase in labeling in the 3xMetRS* mouse line. Scale bar 100 µm. (**B**) Quantification of the experiments shown in A, **** p<0.0001. (**C**) Quantification of ANL incorporation in primary cortical neurons from Nex-Cre:3xMetRS* +/- anisomycin showing the specificity of the labeling for ANL incorporated into proteins, **** p<0.0001. (**D**) GFP expression in Nex-Cre:1xMetRS* and Nex-Cre:3xMetRS* quantified from cultured cells by confocal microscopy, p<0.0001 (**E**) Representative Western Blot showing BONCAT for ANL labeling in cortical primary neurons from Nex-Cre:1xMetRS* and Nex-Cre:3xMetRS* mice +/- ANL. (**F**) Graph displaying the quantification of the experiments shown in E, * p=0.0314, ** p=0.0011. (**G**) Representative confocal images of FUNCAT in acute slices from Nex-Cre:1xMetRS* and Nex-Cre:3xMetRS* mice labeled with ANL for 2 h, showing newly synthesized proteins in the excitatory neurons of the hippocampus. Scale bar 250 µm. (**H**) Quantification of ANL incorporation in primary cortical neurons from Nex-Cre:3xMetRS* and Nex-Cre:6xMetRS* showing increased ANL labeling, *** p=0.0010, * p=0.270. (**I**) Quantification of puromycin incorporation in primary cortical neurons from heterozygous or homozygous 1xMetRS* and 3xMetRS* showing similar incorporation of puromycin +/- Cre. Number of quantified neurons: 1x Cre 979, 2x Cre 428, 3x Cre 286, 6x Cre 286,1x Control 171, 2x Control 951, 3x Control 690, 6x Control 337, 1x Cre no puro 134, 2x Cre no puro 108, 2x Control no puro 163, 6x Control no puro 231.

To examine how the number of allele copies influences ANL labeling, we prepared primary cortical cultures from offspring of crosses leading to different genotypes and compared Nex-Cre::3xMetRS* and Nex-Cre::6xMetRS* with their littermates that did not express Cre. We observed only a slight increase in ANL labeling in 6xMetRS* versus 3xMetRS* in cultured neurons (Fig. 2, H), suggesting that the maximal labeling may plateau under our conditions. In contrast to the copy number dependence of the ANL labeling, we found, as expected, that total protein synthesis levels, as assessed by puromycylation (Schmidt *et al*, 2009), were not different (Fig. 2, I). Thus, ANL incorporation depends, as expected, on Cre expression and the rank order of ANL incorporation scales with the number of encoded MetRS* copies. The scaling between ANL incorporation and encoded MetRS* copy number, however, was not linear and could be influenced by additional factors. For example, there could be a difference in mRNA abundance of the transcripts transcribed from the 1xMetRS* and 3xMetRS* alleles. Since the large expression cassette in the 3xMetRS* line (3.6 kb for 1xMetRS* and 8.6 kb for 3xMetRS*) was expected to be transcribed and translated poorly, we added a WPRE stabilization element to this construct to ensure expression. As a result, the 3xMetRS* cassette expression may be higher. To understand if mRNA abundance contributes to the superior performance of the 3xMetRS* line and the difference in GFP protein expression, we conducted FISH experiments using probes to detect *gfp mRNA,* cell type marker mRNAs (*gad2, camk2a, bdnf*) and the MetRS mRNA *mars1* (Perez *et al*, 2021a). Since only excitatory neurons are expected to express the mutant MetRS* in the crosses with Nex-Cre, we reasoned that inhibitory neurons (*gad2+*) serve as controls in these cultures (Supp Fig. 1A,C-E). Therefore, these assays allowed us, in addition, to address the level of *mars1\** expression in relation to the wild type transcript. Indeed, consistent with the observed higher GFP protein expression mRNA expression of the inserted cassette in cultures was higher in the 3xMetRS* mouse compared to the1xMetRS* as judged by *gfp* FISH signal (Supp Fig. 1B) suggesting stabilization of the large transcript.

The *mars1* FISH signal also revealed a substantial overexpression of the mRNA for MetRS* in the 3xMetRS* compared to the1xMetRS* mouse (Supp Fig 1B, C-E). Although the *mars1* FISH probe does not discriminate between the mutant and wild-type sequence, we addressed the relative expression levels of mRNA for wildtype MetRS and mutant MetRS* by performing dual color FISH labeling in various genotypes comparing distinct cell types (*gfp*+, *gad2*+, *camk2a*+, *bdnf*+) within the cultures from Cre-positive and Cre-negative animals (Supp Fig 1C-E). In the absence of Cre, *mars1* levels were comparable between excitatory and inhibitory cells, reflecting the levels of the wild type enzyme mRNA. In the Nex-Cre::3xMetRS* cultures the signal was significantly higher in excitatory cells (*camk2a*+, *bdnf*+) than in inhibitory cells (*gad*+), as expected. In the Nex-Cre::1xMetRS* cultures overexpression of *mars1* in excitatory neurons was substantially lower and not significantly higher than the wild type enzyme mRNA (Supp Fig. 1B, E). We obtained similar results from FISH experiments on brain slices, where we analyzed the hippocampal CA1 region (Supp Fig. 1F). Taken together, the *in vitro* experiments demonstrated a significantly increased Cre-dependent labeling efficiency of the Cre-defined cell type in the 3xMetRS* line.

### Methionine competition in 3xMetRS* versus 1xMetRS* *in vitro*

We reasoned that increasing the expression of MetRS* would favor the incorporation of ANL over methionine. To address this, we compared ANL incorporation in Nex-Cre::1xMetRS* and Nex-Cre::3xMetRS* neurons in culture media supplemented with methionine or not. Indeed, ANL incorporation in Nex-Cre::3xMetRS* neurons was not only more pronounced than in Nex-Cre::1xMetRS* cultures; it was also not significantly inhibited by the presence of methionine in the growth medium, as observed in Nex-Cre::1xMetRS* neurons (Fig. 3 and Supp Fig. 2). While a slightly reduced (but non-significant) ANL uptake is still visible after 1 hour of ANL incubation in Nex-Cre::3xMetRS* neurons, no difference is observed with 2 hour labeling (Fig. 3C, D). This trend likely reflects the initial presence of methionine-loaded tRNA prior to the start of incubation. In contrast, Nex-Cre::1xMetRS* neurons exhibit strong methionine competition at both time points (Fig. 3A, B).

**Figure 3.**
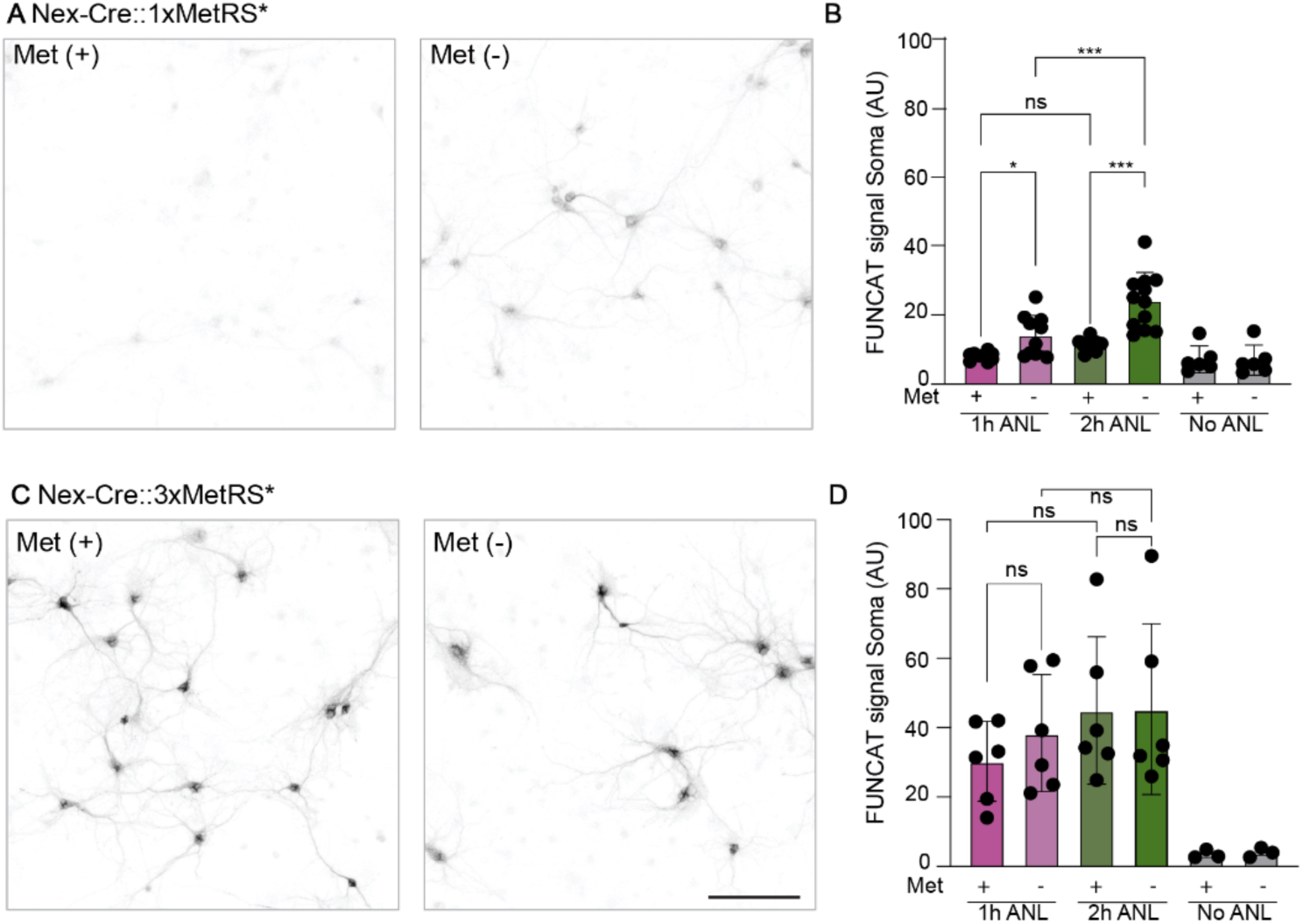
Methionine competition with ANL. (**A, C**) FUNCAT in primary neurons from cortices of Nex-Cre::1xMetRS* mice (A) or Nex-Cre::3xMetRS* mice (C) labeled with ANL in the presence (met+) or absence (met-) of methionine in the culture media. (**B**, **D**), Quantification of the experiments depicted in A and C, respectively* p=0.0437, *** p=0.0001, **** p<0.0001 (2h +/- met). Each dot represents the average of single-cell quantifications for one dish of at least 3 independent experiments. (Number of neurons quantified: Nex-Cre::1xMetRS*; 1h +met (2085), 1h-met (1496), 2h +met (1904), 2h-met (1651), no ANL +met (1189), no ANL-met (1236). Nex-Cre::3xMetRS*; 1h +met (814), 1h-met (637), 2h +met (558), 2h-met (595), no ANL +met (520), no ANL-met (460)). Bars represent mean ±SD. Scale bar 100 µM.

### ANL *in vivo* labeling in 2xMetRS* and 3xMetRS* mouse line

The above enhanced *in vitro* labeling in Nex-Cre::3xMetRS* prompted us to examine *in vivo* ANL labeling. We administered ANL via a single intraperitoneal injection and compared labeling from 0.5 to 6 hours and compared the “best” first-generation mouse line (homozygous, Nex-Cre::2xMetRS*), to heterozygous mice from the new generation line (Nex-Cre::3xMetRS*) (Fig. 4A). As expected from earlier studies, no BONCAT signal was detectable after short labeling periods in cortex samples from the Nex-Cre::2xMetRS* mice; in contrast, protein synthesis was clearly detected from Nex-Cre::3xMetRS* mice with labeling times of 1 hour or longer (Fig. 4B, C). This finding demonstrates that ANL incorporation is achieved much faster in the new mouse model, presumably due to enhanced efficiency of ANL loading and reduced methionine competition (Fig. 3 and Supp Fig. 2).

**Figure 4.**
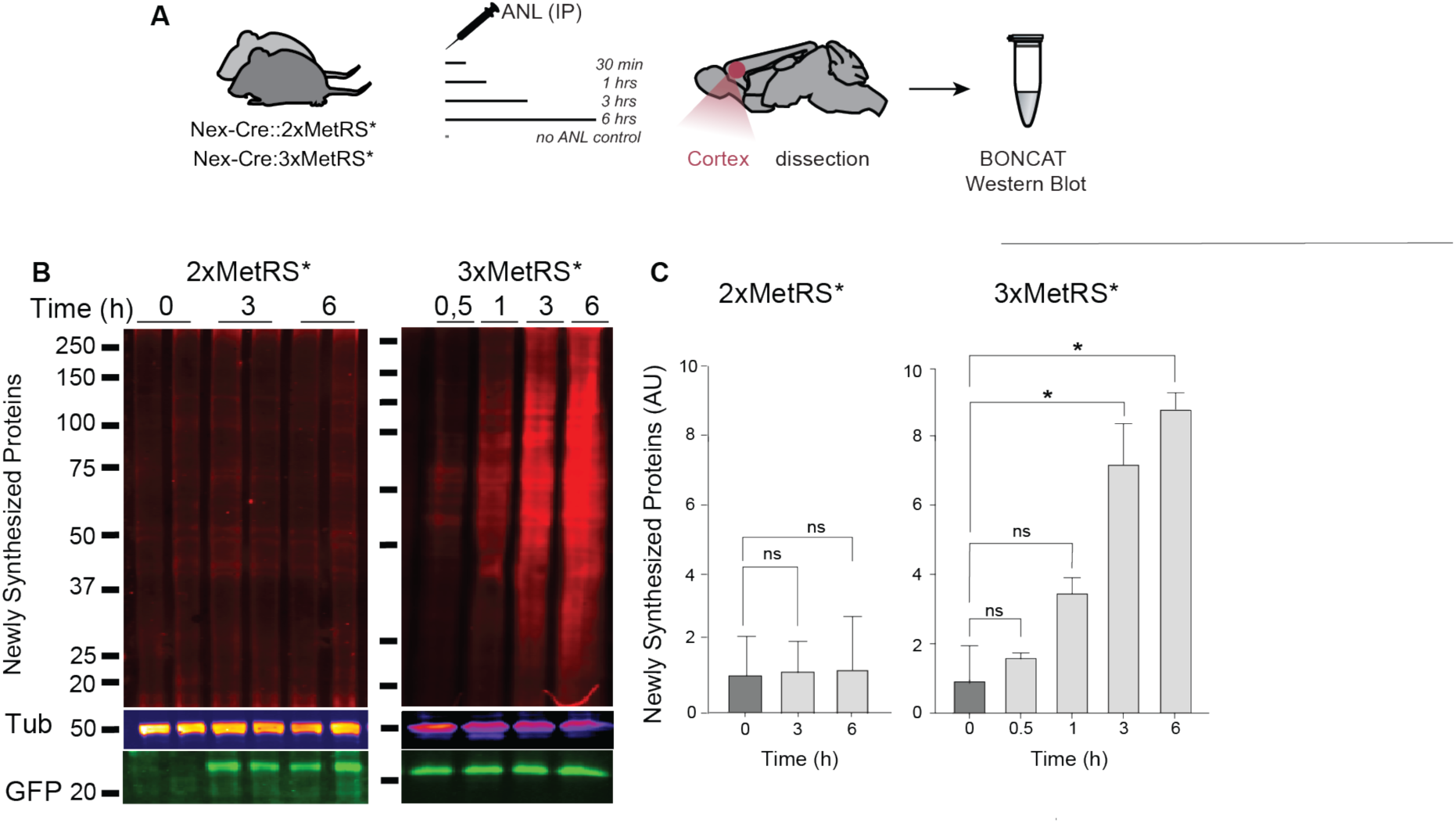
*In vivo* ANL labeling efficiency in 2xMetRS* vs. 3xMetRS*. (**A**) Experimental outline: Nex-Cre::2xMetRS* and Nex-Cre::3xMetRS* cortices were collected 0.5 hours, 1 hour, 3 hours, or 6 hours after a single ANL IP injection. (**B**) Western blot of newly synthesized proteins labeled with ANL and detected by BONCAT. Tubulin is used as a control for protein loading, and GFP expression serves as a proxy for MetRS* expression. (**C**) Bar graph showing the quantification of newly synthesized proteins shown in B, * p=0.0307 (0 / 3h), * p=0.0106 (0 / 6h). Error bars represent SD.

### ANL *in vivo* labeling in the 3xMetRS* and 6xMetRS* mouse line

To study the effect of allele copy number in the new line, we next compared heterozygous (Nex-Cre::3xMetRS*) and homozygous mice (Nex-Cre::6xMetRS*). As before, we analyzed the BONCAT signal from tissue labeled with ANL for short periods after a single IP injection to compare protein labeling during the exponential phase. Hippocampal (HC) tissue was collected at 1 hour, 3 hours, and 6 hours (Fig. 5A) after a single ANL injection, and protein synthesis was detected with increasing levels in both heterozygous and homozygous animals (Fig. 5B, C). While GFP expression was—as expected—doubled in homozygous animals compared to heterozygous ones, the BONCAT signal was higher as a trend at each of the studied time points but slightly less than double (Fig. 5B, C). This fits with the findings from the *in vitro* experiments and suggests an initial saturation of tRNA loading. Thus, excitingly, even though labeling is slightly better in homozygous mice, it is still possible to use heterozygous animals for labeling. The protein synthesis signal is also detectable by FUNCAT in hippocampal sections from heterozygous animals after 3 hours of labeling (Fig. 5D). We note that the ability to use heterozygous animals represents a significant reduction in the breeding burden of mice, especially for experiments addressing cell type-specific proteomes in disease or mutation conditions.

**Figure 5.**
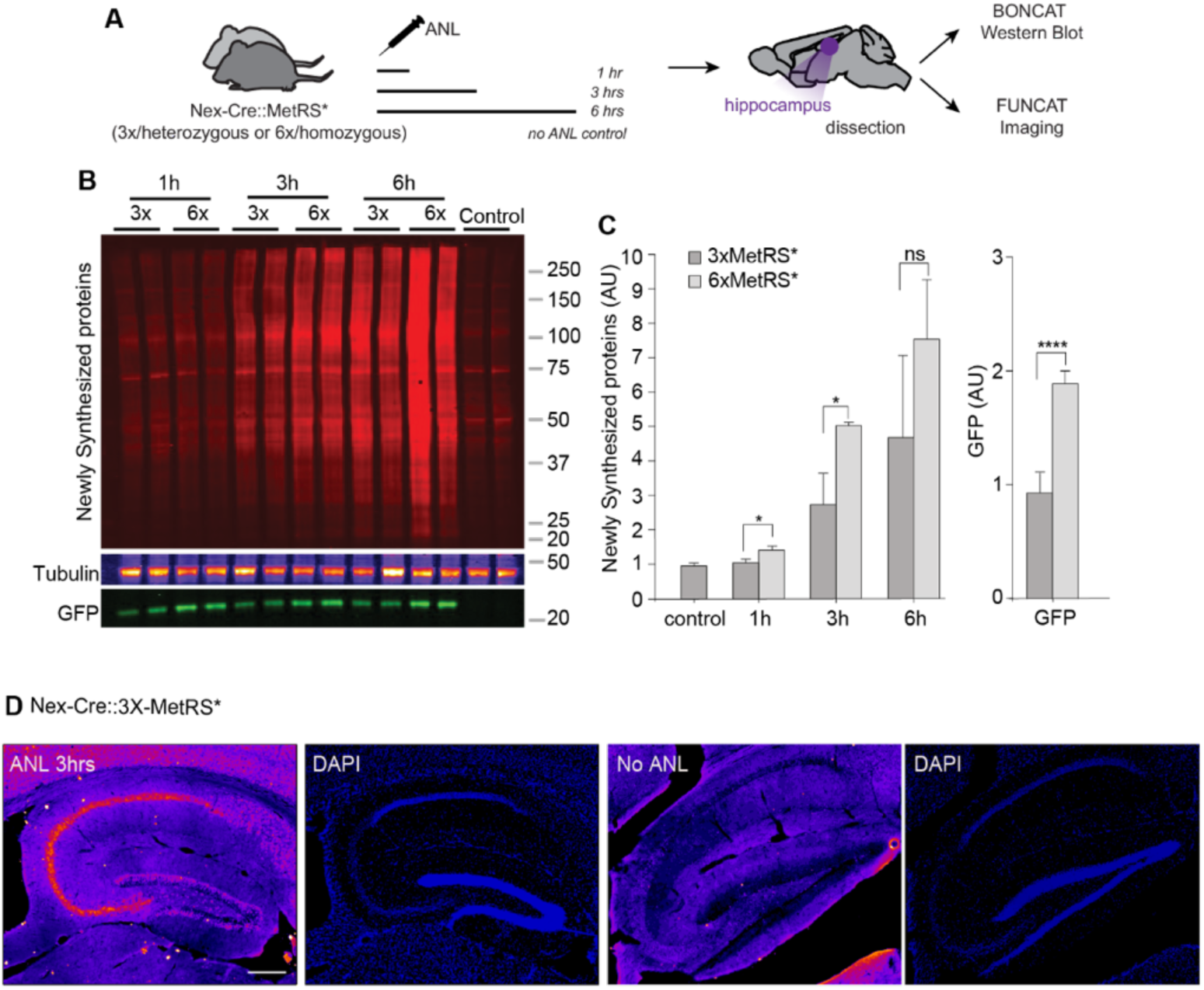
*In vivo* ANL labeling efficiency in 3xMetRS* vs. 6xMetRS*. (**A**) Experimental outline: ANL was administered to Nex-Cre::3xMetRS* and Nex-Cre::6xMetRS* mice via a single IP injection and hippocampal tissue was collected after 1 hour, 3 hours, and 6 hours. (**B**) Western blot of newly synthesized proteins labeled with ANL and detected by BONCAT. Tubulin was used as a loading control, and GFP serves as a proxy for MetRS* expression. (**C**) Bar graph showing the quantification of newly synthesized proteins, * p=0.0382, ** p=0.0030, **** p<0.0001. (**D**) Quantification of GFP expression shown in B. (**E**) FUNCAT images of Nex-Cre::3xMetRS* and negative control brain sections collected 3 hours after IP injection, showing specific ANL labeling in the excitatory neurons of the hippocampus. Scale = 200 µm. Error bars represent SD.

### Identification of newly synthesized proteins via mass spectrometry (MS) with temporal control

To demonstrate the feasibility of identifying newly synthesized proteins using the 3xMetRS* mouse line, we identified nascent proteins from cortical excitatory neurons labeled for 3 and 6 hours (Fig. 6A and Supp Fig. 3A). The cortices from individual mice served as replicates. We analyzed the proteins detected exclusively or enriched, with at least a 3-fold increase (FDR < 0.01) compared to wild type mice. We quantified more than 8,000 proteins in each dataset, of which 3,963 and 5,876 proteins were enriched or exclusive at 3 hours and 6 hours of labeling (Fig. 6B, C, and Supp Fig. 3B-F). Additionally, at 6 hrs, the overall intensity of the identified proteins significantly increased (Fig. 6D). To confirm whether we could detect different protein pools by simply changing the ANL labeling period, and considering that newly synthesized proteins typically have shorter half-lives and are more dynamic (Ross *et al*, 2021; Tong *et al*, 2020), we compared the 3- and 6-hour proteomes to previously published protein half-life datasets in a density plot (Fig. 6E and Supp Fig. 3G) (Dorrbaum *et al*, 2018; Fornasiero *et al*, 2018). We found that the identified proteins at 3 hours were shifted toward shorter (6.25 days, median) half-lives, while those at 6 hours are shifted toward longer (6.95 days, median) half-lives, whereas the overall median half-life of the cortical proteins used as a background is 8.09 days (Fig. 6E). As such, by labeling with ANL for different durations, we alter the sensitivity for detecting proteins with different half-lives. Next, we compared the identified proteomes with each other (Fig. 6F-H and Supp Fig. 3H) and found 3,823 common proteins between both labeling times, with 140 and 2,053 proteins exclusively identified at 3 hours and 6 hours, respectively. Looking into the common proteins in both datasets, we found 758 proteins significantly more abundant at 6 hours, while only 2 proteins were found to have higher intensities at 3 hours (at least 1.5-fold, FDR < 0.01).

**Figure 6.**
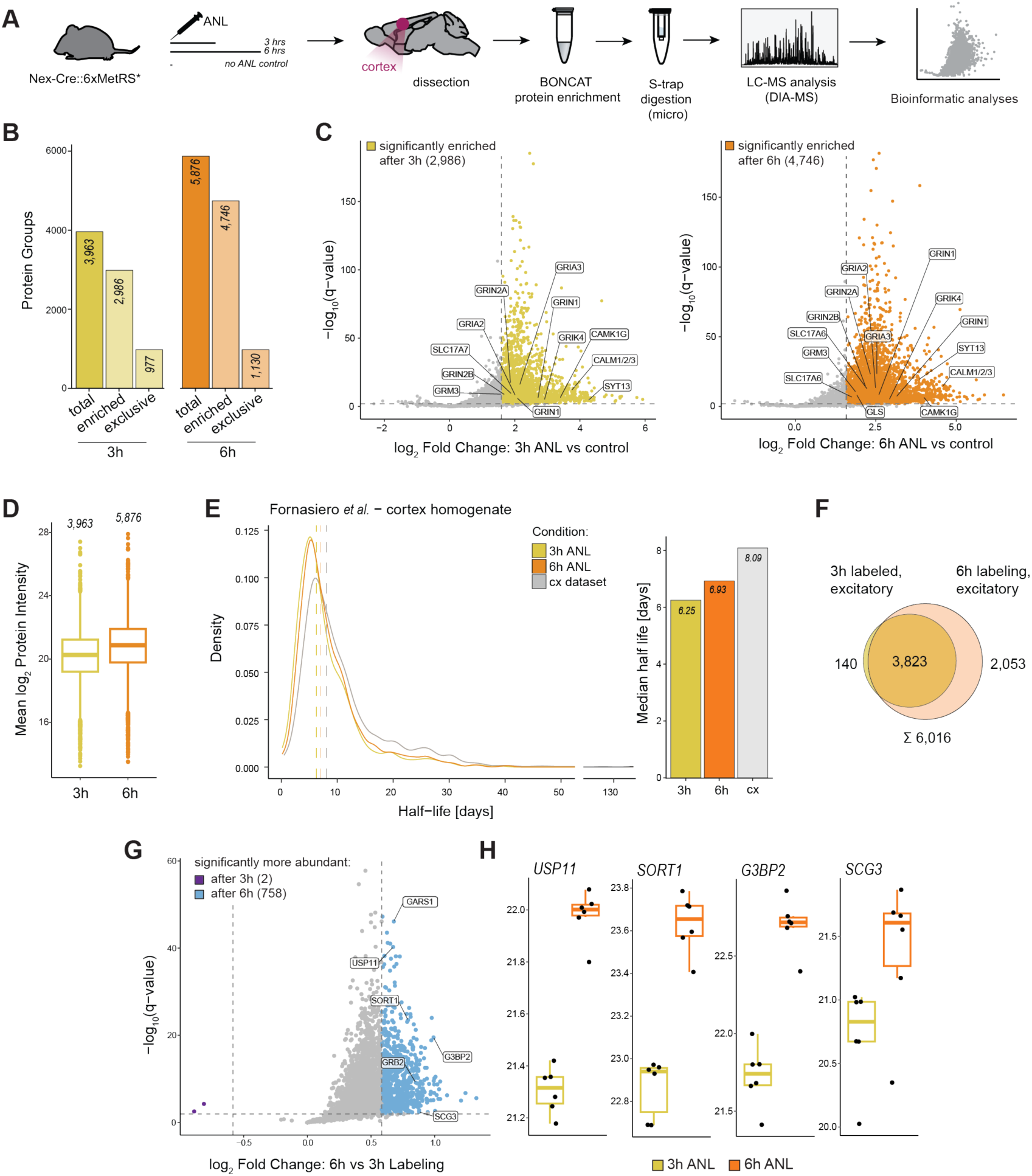
Excitatory neuronal protein identification after 3 and 6 hours of *in vivo* ANL labeling. (**A**) Workflow for labeling and purification of excitatory neuronal proteins. *In vivo* protein labeling was performed by intraperitoneal injection of ANL in Nex-Cre::6xMetRS* mice. Cortex tissue was collected after 3 hours or 6 hours post-injection (n = 6) or no injection (no ANL control, n = 2). Proteins containing ANL were clicked, purified (BONCAT), and quantified by LC-MS. (**B**) Bar plot shows protein group quantifications of the excitatory neuronal proteome after 3 hours and 6 hours of labeling. Excitatory proteomes were defined by either exclusive identification in ANL-labeled samples or by significant enrichment over the unlabeled controls. See also Supp Fig. 3A for method details. (**C**) Volcano plot comparing protein abundance of shared proteins in the 3-hour (left) and 6-hour (right) samples to the unlabeled control (peptide-level linear mixed effects model, Benjamini-Hochberg p-value correction). Significantly enriched protein groups (80% valid value in labeled condition, at least 3-fold enrichment, FDR < 0.01) for either time point are highlighted in color (3 hours – yellow; 6 hours – orange). Selected candidate proteins of excitatory synapses are labeled using their gene symbols. (**D**) Box plot displays the mean log2-scaled intensities of the newly synthesized excitatory proteins after 3 hours and 6 hours of labeling. Significant increases in protein intensity were observed with prolonged labeling duration (two-sided, unpaired Welch’s t-test, p-value = 2.2e-16; 1.54-fold enrichment). Boxes indicate the median, first, and third quartiles, while whiskers extend to 1.5 × IQR. (**E**) The density plot (left) shows the protein half-life distribution. Excitatory neuronal proteins identified at each time point were matched to a previously published database for protein half-lives in the brain cortex (Fornasiero *et al*., 2018) and are highlighted by color; full cortex data is depicted in gray. Vertical lines in the diagram and bar plot (right) highlight the increase in median half-lives of the matched excitatory proteins after 3 hours (6.25 days) and 6 hours (6.93 days) of labeling compared to the median half-lives of all cortical proteins (8.09 days). (**F**) The Venn diagram compares the excitatory proteomes of the two time points, indicating high overlap between them. (**G**) The volcano plot compares the shared excitatory protein groups quantified after 3 hours and 6 hours of labeling (peptide-level linear mixed effects model, Benjamini-Hochberg p-value correction). Significantly more abundant proteins (at least 1.5-fold enrichment, FDR < 0.01) at either time point are highlighted by color (3 hours –purple; 6 hours – blue). H, Box plots display selected candidate proteins showing significantly higher abundance after 3 hours (ARC, EGR1) or 6 hours of labeling (USP11, SORT1, G3BP2, SCG3). Boxes indicate the median, first, and third quartiles, while whiskers extend to 1.5 × IQR.

### A cell type-specific proteome from a low abundance neuronal population: dopaminergic neurons

Given the increased labeling sensitivity in the 3xMetRS* mouse line, we next tested whether we could identify proteomes from neurons that account for low numbers in a given brain region. As a proof of principle, we chose the dopaminergic neurons from the substantia nigra-SN- and the olfactory bulb-OB-. In the SN, there are approximately 14,000 dopaminergic neurons in each mouse (Ip *et al*, 2017), and in the OB, the DA neurons represent 6% of the OB cells, totaling about 89,000 cells (Parrish-Aungst *et al*, 2007). By pooling dissected SN from several animals (5 per replicate), we were able to detect nascent protein labeling by BONCAT. However, the amount of purified protein was not sufficient for accurate protein identification following MS. Thus we next focussed on the OB and identified, for the first time, the DA neuron-OB proteome, using DAT-Cre::6xMetRS* animals labeled with ANL for 2 weeks (Fig. 7A, B, and Supp Fig. 4A).

**Figure 7.**
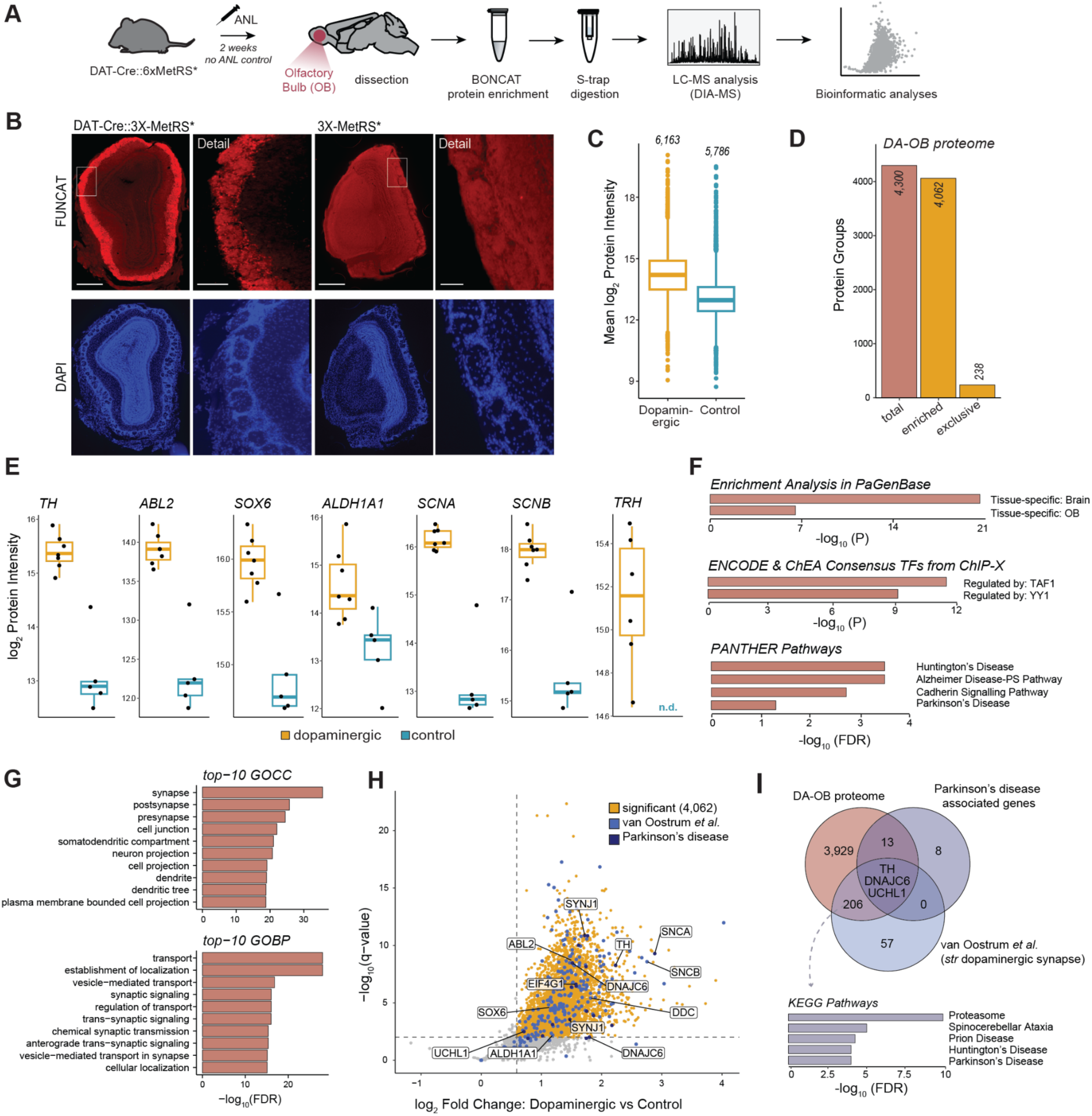
Olfactory bulb dopaminergic proteome identification. (**A**) Workflow for labeling and purification of dopaminergic neuronal proteins. *In vivo* protein labeling was performed by daily intraperitoneal injection of ANL in DAT-Cre::6xMetRS* mice. Olfactory bulb tissue was collected after two weeks for labeled samples (n = 7) or no-ANL control samples (n = 5). Proteins containing ANL were clicked, purified (BONCAT), and quantified by LC-MS. (**B**) Confocal microscopy images showing the ANL-labeled proteins by FUNCAT in the DA neurons of the OB. (**C**) Boxplot of overall protein group intensities (log2) of the labeled condition and unlabeled control, merged by mean values, showing that most proteins identified in the OB-DA neuron samples are enriched compared to the control samples. Boxes specify the median, first and third quartiles, and whiskers extend to 1.5 × IQR. (**D**) Barplot showing protein group identifications of the dopaminergic neuronal proteins quantified from the olfactory bulb. The OB-DA neuron proteome was defined by either exclusive identification in ANL-labeled samples or by significant enrichment over the unlabeled controls. See also Fig. S4A for method details. (**E**) Boxplots display brain disease-associated and/or previously reported dopaminergic candidate proteins showing significant enrichment in the labeled over the unlabeled control samples (TH, ABL2, SOX6, ALDH1A1, SCNA/B) or exclusive detection in the labeled conditions (TRH). Boxes specify the median, first and third quartiles, and whiskers extend to 1.5 × IQR. (**F**) Overrepresentation analysis showing enriched terms for the dopaminergic proteome according to annotation in PaGenBase, transcription factor ChIP-seq data from ENCODE (assessed via Enrichr, OB background), and annotation of cellular pathways by PANTHER (OB background). (**G**) Gene ontology overrepresentation analyses display the top 10 terms for the dopaminergic proteome according to the annotation of cellular compartment (GOCC) or biological process (GOBP) using the OB proteome as background (FDR < 0.05). (**H**) Volcano plot comparing shared proteins quantified in labeled samples to the unlabeled controls (peptide-level linear mixed effect model, Benjamini-Hochberg p-value correction). Significantly enriched protein groups (80% valid value in labeled condition, at least 1.5-fold enrichment, FDR < 0.01) are highlighted in yellow. Furthermore, color indicates proteins matching the dopaminergic proteome of striatal synapses or Parkinson-associated genes (UniProt KW: KW-0907). Selected candidate proteins of dopaminergic neurons are labeled using their gene symbols. (**I**) Venn diagram highlighting the overlap of the total dopaminergic proteome with dopaminergic proteins of striatal synapses or Parkinson-associated genes (UniProt KW: KW-0907) at the level of gene symbols. Genes shared between the OB-DA neuron proteome and the striatal dopaminergic synapse proteome from van Oostrum et al. show an enrichment in proteasome and neurodegenerative disease proteins (KEGG pathways, FDR < 0.05). Scale bar 200 uM.

We identified 4,300 proteins derived from the DA neurons of the OB (Fig. 7C, D, and Fig. S4B-E), of which 4,062 are enriched by at least 1.5-fold in the OB DA neuron dataset (when compared to control) and 238 were exclusive to the labeled conditions (Fig. 7D, E). We used several databases, such as Metascape (Pan *et al*, 2013; Zhou *et al*, 2019), to confirm that the identified proteins in our OB-DA neurons are specific to the OB and brain tissue. Overrepresentation analysis of the OB-DA proteome compared to a full OB proteome (used as background, (Sharma *et al*, 2015)) in the Enrichr database (Chen *et al*, 2013; Kuleshov *et al*, 2016; Xie *et al*, 2021) with transcription factor ChIP-seq data from ENCODE, indicated increased expression of genes regulated by TAF1 and YY1 (Ying Yang 1) (Fig. 7F). We also performed an overrepresentation analysis (using the PANTHER pathway database; (Mi *et al*, 2007)) and found terms associated with three neurodegenerative diseases: Alzheimer’s disease, Huntington’s disease, and Parkinson’s disease (PD) (Fig. 7F). Additionally, as expected, there was pronounced overrepresentation of gene ontology terms related to neurons and synapses (Fig. 7G). Next, we compared the DA-OB proteome with the proteome of DA neuronal synaptosomes isolated from the striatum and detected several common proteins (209) (van Oostrum *et al*, 2023). These common proteins show significant enrichment in terms related to the proteasome, which plays an important role in PD and other proteinopathies (Fig. 6H, I).

## Discussion

Understanding how protein homeostasis works *in vivo* is essential for fully grasping the internal processes of cells and organisms. Complex organisms comprise hundreds of distinct cell types, each responding differently to identical physiological or pathological challenges due to their specialization. The method described here uses the expression of a mutant MetRS*, which allows for the incorporation of a bioorthogonal analogue of methionine (ANL) into proteins, enabling protein purification and increasing detection sensitivity by MS. In the present work, we compared a first-generation mouse line expressing one copy of the MetRS* enzyme per allele (1xMetRS*) with a second generation expressing three copies of MetRS* (3xMetRS*) per allele. In both mouse lines, MetRS* expression is Cre recombinase-dependent. With the 3xMetRS* line, we increased ANL labeling efficiency, reduced the time of labeling to study protein synthesis over a shorter time frame, and identified a cellular proteome in a very small population of dopaminergic neurons.

One drawback of metabolic labeling with amino acid analogs is competition with the corresponding natural amino acid which can also be charged by the MetRS*. To achieve effective labeling, experiments with the first-generation MetRS* line—both *in vitro* and *in vivo*—were necessarily conducted under methionine-free or -reduced conditions to prevent competition between methionine and ANL. Here we obtained strong labeling without altering physiological methionine concentrations. We demonstrated that cell type-specific protein identification is robust just 3 hours after a single ANL IP injection. Additionally, when comparing the proteins identified after 3 hours with those identified after 6 hours, we identified protein pools with different half-lives, highlighting the potential of the 3xMetRS* system to study newly synthesized proteins that are missed with the 1xMetRS* system The growing interest in the study of proteins and their behavior has led to the continuous development of methods various techniques (Alvarez-Castelao *et al*., 2017; Ignacio *et al*, 2023; Kim *et al*, 2016; Kubitz *et al*, 2022; McClatchy *et al*, 2015; Ong & Mann, 2006; Schmidt *et al*., 2009; Villalobos-Cantor *et al*, 2023). Despite this progress, BONCAT (Dieterich *et al*., 2006), and specifically ANL labeling remains the most versatile method for studying protein homeostasis *in vivo*. The strength of this system lies in its cell type specificity, allowing for *in situ* visualization of proteins, and its reliance on metabolic labeling. Depending on the labeling duration, ANL can be utilized to investigate different aspects of protein homeostasis: 1) studying protein synthesis by identifying proteins after brief labeling periods; 2) examining protein steady state or proteomes by identifying proteins after longer labeling periods that exceed the average half-lives of the proteins in the studied tissue or cell type; and 3) analyzing protein degradation by ceasing ANL administration after an initial labeling (t=0) and tracking the decrease of labeled proteins over time. Additionally, the possibility to identify the cell type-specific proteomes from a single animal allows one to correlate the (behavioral or disease) phenotype of individual mice with their respective proteomes. Other methods, such as puromycylation, are extensively used for studying protein synthesis due to their high sensitivity. Puromycin is incorporated at any location within the translating protein. This method allows for the investigation of protein synthesis after short labeling periods (e.g., 5-10 minutes). However, prolonged labeling with puromycin can lead to cellular toxicity and the method is not cell type-specific, though recent developments have enabled puromycin-resistance or activation in identified cell types (Cabrera-Cabrera *et al*, 2023; Villalobos-Cantor *et al*., 2023).Another useful approach is TurboID and its variations. This method relies on biotin binding to proteins in close proximity through an enzymatic reaction. Like puromycylation, BioID is highly efficient, allowing for brief labeling periods. However, it is primarily used to study local proteomes or protein-protein interactions and is not suitable for studying protein synthesis, as it does not involve metabolic labeling.

The 3xMetRS* line also enables the identification of proteomes from low-abundance neuronal populations, such as olfactory bulb dopaminergic neurons (representing just 6% of the cells in the bulb). Regrettably, we were, as yet, unable to purify and identify the proteome from the dopaminergic neurons of the substantia nigra using a similar number of neurons as starting material. This may indicate that the protein synthesis rate of dopaminergic neurons in the bulb is higher, likely influenced by the retained capacity of the olfactory bulb dopaminergic neurons for postnatal proliferation. Indeed, this feature positions the olfactory bulb dopaminergic neurons as potential instruments for restorative treatments for Parkinson’s Disease, and they could even serve as an autologous source for transplantable dopaminergic neurons (Baker *et al*, 2001). In addition, dopaminergic neurons in the olfactory bulb can modulate odor processing and discrimination (Banerjee *et al*, 2015; Escanilla *et al*, 2009). One of the main non-motor symptoms affecting up to 90% of Parkinson’s Disease cases is smell dysfunction, which can occur years before motor symptoms manifest (Doty, 2012). Hyposmia is considered a potential biomarker for Parkinson’s Disease; however, the underlying pathways and their connection to substantia nigra degeneration are poorly understood. Here, we identified, for the first time, the proteome of dopaminergic neurons from the olfactory bulb, showing enrichment in the transcription factors TAF1 and YY1, as well as in proteins associated with the proteasome, all of which are closely related to Parkinson’s Disease and other neurodegenerative diseases. Interestingly, hemizygous mutations in TAF1 have been associated with X-linked dystonia and Parkinson’s disease in humans, as well as with other disorders such as global developmental delay and intellectual disability (Makino *et al*, 2007; O’Rawe *et al*, 2015; Zeng *et al*, 2022). YY1 miss-regulation is closely related to a myriad of brain disorders, including neurodegenerative diseases such as AD and PD (Pabian-Jewula *et al*, 2022; Tiwari & Pal, 2017).

## Methods

### Transgenic animals

For the new mouse line the cassette —STOPFLOX-3x(MetRS*-2A)-GFP— is expressed under the CAG actin-derived promoter. The line was custom made by knock-in at the mouse ROSA26 locus using recombination-mediated cassette exchange (RMCE) in embryonic stem cells (Taconic GmbH). For expression of the cassette in excitatory neurons, homozygous floxed 3xMetRS* (available under request to BAC) animals or 1xMetRS* animals (JAX #028071) were crossed with homozygous Nex-Cre mice (kindly provided by K.A. Nave, (Goebbels *et al*., 2006) or DAT-Cre mice (JAX #006660), (Backman *et al*, 2006). Genotyping of the 3xMetRS* and wild type alleles was conducted by PCR in separate reactions with KAPA2G Fast (HotStart) Genotyping Mix (wt allele: Primers CTC TTC CCT CGT GAT CTG CAA CTC C, CAT GTC TTT AAT CTA CCT CGA TGG, internal control primers:GAG ACT CTG GCT ACT CAT CC, CCT TCA GCA AGA GCT GGG GAC knock-in allele primers: ACT GGC TAC CTT GTC TGA GGA G, CAT GGA TGG CAT GGT ACT TG, 94°C 1min, 35x (94°C 30sec, 62°C 30sec, 72°C 1min) 72°C 10min, 4°C). All experiments involving animals were performed with permission from the local government offices in Germany (RP Darmstadt; protocol V54-19c20/15-F126/2002) or Spain (Committee of Animal Experiments at UCM and Environmental Counseling of the Comunidad de Madrid, protocol number: PROEX 005.0/21). These experiments comply with the German Animal Welfare Law, Max Planck Society rules, Spanish regulations, and follow EU guidelines for animal welfare. Mice were housed in accordance with EU Directive 2010/63/ EU Annex III and sacrificed using methods in accordance with EU Directive 2010/63/ EU Annex IV.

### Antibodies

The following antibodies were used for immunofluorescence labeling-IF- and/or immunoblotting-IB-at the indicated dilutions. Primary antibodies: rabbit anti-biotin (IB, 1:1000-, Cell Signaling 5597), rabbit anti-GFP (IF in cells 1:500, abcam ab6556) or chicken anti-GFP (IF, 1:500; IB: 1:1,000; Aves), chicken anti-GFP (IB, IF 1:500, abcam ab13970), guinea pig anti-MAP2 (IF in cells 1:1,000, Synaptic Systems 188004), mouse anti-Puromycin (IF in cells 1:3500, Kerafast EQ0001). Secondary antibodies: goat anti-mouse, anti-rabbit or anti-chicken IRdye680 or IRdye800 (IB, 1:10,000, Licor), goat anti-guinea pig-Dylight405 (IF in cells 1:1000, Jackson ImmunoResearch 106-475-003), goat anti-rabbit Alexa488 (IF in cells, 1:1000, Thermo Fisher Scientific A11008), goat anti-chicken Alexa488 (IF in slices, 1:1000, Thermo Fisher Scientific A11099), goat anti-mouse Alexa546 (IF in cells 1:1000, Thermo Fisher Scientific A11030). In slice IF experiments, DAPI was added to the secondary antibody solutions or to wash solutions where appropriate.

### Primary cultures

Primary neuronal cultures were prepared from pooled litters or single animals of either sex from crosses of homozygous Nex-Cre females with homozygous 1x/3x MetRS* males or double heterozygous Nex-Cre::1x/3xMetRS* females with the respective homozygous 1x/3xMetRS* males and maintained as previously described (Aakalu *et al*, 2001). Briefly, cortices from newborn (P0/P1) transgenic mice, were dissected and dissociated with papain (Sigma P3125), washed, triturated and plated at a density of 30.000 cells/coverslip on poly-D-lysine coated MatTek glass bottom dishes (MatTek P35G-1.5-14C) for imaging experiments or 3 Mio cells on poly-D-lysine-coated 60 mm cell culture dishes for BONCAT. Neurons were maintained at 37°C and 5% CO_2_ in NGM (Neurobasal-A plus B27 and GlutaMAX) supplemented with glia- and cortex-conditioned NGM. Experiments were performed at DIV 7-14.

### BONCAT in cultured neurons

Growth medium was removed and replaced with NGM without methionine (Gibco, custom preparation) supplemented with 4 mM ANL (Iris Biotech HAA1625 or Alvarez-Castelao et al. 2017) or 4 mM methionine. Following metabolic labeling for 2 hr cultured neurons were washed two times with D-PBS (Gibco) on ice and harvested in D-PBS by scraping and a 40-second spin-down in a small bench top mini centrifuge. The supernatant was removed and the pellet immediately frozen on dry ice and stored at −80°C until use. Lysates were prepared by resuspension in 60 µl PBS pH7.8 supplemented with 0.4 % (w/v) Triton X-100 and 0.4 % (w/v) SDS, along with protease inhibitor (PI, 1:1000 dilution of protease inhibitor cocktail 3 w/o EDTA, Calbiochem) and benzonase (Sigma, 1:1,000) at room temperature and incubation at 75 °C for 10 min. Lysates were then cleared by centrifugation for 10 min, 13.000 x g at 15 °C and protein measured by BCA assay (Invitrogen). Click chemistry was performed by adding to 100 μL PBS pH 7.8 (with Protease Inhibitor 1:4000) 20 µl lysate with 100 μg protein, 500 μM triazole ligand Tris[(1-benzyl-1H-1,2,3-triazol-4-yl)methyl]amin (TBTA, Thermo Fisher Scientific, 454531000), 62.5 μM biotin-alkyne tag (Thermo Fisher Scientific, B10185) and 83 μg/mL CuBr (prepared by dilution of a fresh 10 mg/mL solution in DMSO). The reaction was incubated at room temperature overnight in the dark by overhead rotation. Biotinylated proteins were separated by SDS-PAGE, immunoblotted with anti-GFP and anti-biotin antibodies and IRDye secondary antibodies and scanned with a Licor Odyssey fluorescence Imager.

### FUNCAT in cultured neurons

Growth medium was removed and replaced with NGM without methionine (or with methionine in the methionine competition experiments) supplemented with 4 mM ANL or 4 mM methionine as control. Labeling was performed for the specified duration. After metabolic labeling with ANL for the indicated times, cells were washed once fast with medium and were then placed back in medium without ANL for 10 min into the incubator. For protein synthesis inhibition controls, neurons were pre-incubated for 30 min with 40 µM anisomycin and 40 µM anisomycin was also present during the incubation with ANL. Neurons were fixed in PFA-Sucrose (PBS pH7.4 containing 4% Sucrose, 4% PFA, 1mM MgCl_2_ and 0.1mM CaCl_2_) for 20 min at RT, permeabilized with 0.5% Triton in Blocking buffer (BB: 4% goat serum in PBS), blocked for 1 h in BB and equilibrated in PBS pH 7.8. The click reaction was performed for 2 hrs at room temperature using 2 μM of the Alexa647 alkyne (Thermo Fisher Scientific, A10278), in a copper-mediated click reaction with CuSO4, TCEP and the Triazole ligand TBTA in PBS pH7.8. Following the click reaction, neurons were washed extensively and blocked with BB for 1 hour before immunostaining. Primary antibodies (MAP2, GFP) were applied for 1 hour at room temperature in BB, followed by three washes with PBS (pH 7.4). Secondary antibodies were added for 30 minutes, the samples were washed with PBS and water and mounted with AquaPolymount for imaging.

### FUNCAT in acute hippocampal slices

Hippocampal slices (300 µm) were prepared from Nex-Cre::1x/3xMetRS* adult animals. Mice were deeply anesthetized with Isoflurane and decapitated, the brain was quickly removed and put in ice-cold, slushed sucrose-based saline solution oxygenated with 95 % O_2_/5 % CO_2_, cut first along the longitudinal fissure and then cut with an angle of 30 degrees with a vibratome (Leica VT 1200 S, Germany) to transverse slices in ice-cold, oxygenated sucrose saline solution. Slices were recovered for 1 hr submerged at RT in artificial cerebrospinal fluid (ACSF; in mM: NaCl 125, NaHCO_3_ 25, KCl 2.5, NaH_2_PO_4_ 1.25, glucose 10, CaCl_2_ 2, MgCl_2_ 1) oxygenated with a 95 % O_2_/5 % CO_2_ gas mixture. Slices were then transferred to an interface chamber, placed on top of lens cleaning paper and continuously perfused with oxygenated ACSF another hour at 32°C (perfusion rate 2 ml/min). Subsequently slices were labeled for 2 hours with or without 1 mM ANL in ACSF by perfusion. Following ANL incubation the slices were perfused for 10 min with ACSF to complete incorporation, rinsed twice in ACSF and fixed in 4% PFA, 4% Sucrose in PBS for 1 hr at room temperature, then washed in PBS and cryoprotected in 15 % and 30 % Sucrose in PBS overnight. Slices were then resliced to 30 µm with a cryotome and postfixed 10 min with 4% PFA, 4% Sucrose in PBS, washed and permeabilized overnight at 4°C (0.5% Triton X-100 in BB). Samples were washed twice in PBS pH 7.8 and “clicked” for 3 days. The click chemistry reaction was set up by adding the reagents in the following order: TCEP, TBTA, 2 μM of Alexa647 alkyne, CuSO_4_. After extensive washing, slices were incubated overnight at 4 °C with primary antibodies in BB, washed and incubated for 5 hours at room temperature with secondary antibodies in BB. DAPI was added for 5 min in BB. The sections were again washed before mounting on microscope slides. Fluorescence imaging was performed with a LSM780 laser scanning confocal microscope (Zeiss, Zen10 software) using a Plan-Apochromat 20x/NA0.8 M27 objective and appropriate excitation laser lines and spectral detection windows.

### Puromycilation

Neuron cultures from different genotypes were incubated with 3 µM Puromycin in growth medium for 5 min at 37°C, incubation was stopped by two fast washes with prewarmed medium and cells were fixed in PFA-Sucrose for 20 min. The cells were then permeabilized for 15 min with 0.5% Triton X-100 in BB, blocked in BB for 1 hr and immunostained for Puromycin (1 hr anti-Puromycin, 30 min secondary antibody), then blocked again and post-stained for GFP and MAP2 1hr with primary antibodies and 30 min with secondary antibodies and mounted with AquaPolymount for imaging.

### mRNA FISH

For high sensitivity FISH cultured neurons were fixed for 20 min with PFA-Sucrose. ViewRNA FISH (ViewRNA ISH cell assay, Thermo Fisher Scientific) was performed as described previously (Alvarez-Castelao *et al*, 2020) using the manufacturers detergent for 5 min permeabilization and all probe sets (incubation time 3h, Thermo Fisher ViewRNA probe sets: *mars1* type1, cell marker mRNAs type6) and PreAmp/Amp/LP reagents (incubation time 1h, type1-LP550, type6-LP650) at 1:100 dilution following the manufacturer’s instructions. After mRNA FISH labeling cells were subjected to MAP2 and GFP immunostaining, mounted in AquaPolymount (Polysciences) and imaged.

For mRNA FISH on hippocampal tissue slices, mice of the respective genotypes were sacrificed by decapitation under isoflurane anesthesia, the head was quickly dipped into liquid nitrogen and the brain was dissected out and fixed in 4% PFA, 4% Sucrose in PBS pH 7.4 for 1 h at 4°C and 1h at RT and cryopreserved in 15% Sucrose and 30% Sucrose in PBS in overnight steps at 4 °C before cryotome sectioning to 30 µm hippocampal sections. Sections were postfixed for 10 min with above fixation solution and permeabilized for 20 min with the ViewRNA ISH cell assay detergent solution. The ViewRNA assay was carried out essentially as described above but with overnight incubation in the probe set mixture. Label probes coupled to the 550dye were used for assays on tissue slices. Subsequent immunostaining was carried out by overnight incubation in primary antibodies at 4°C and 4hr secondary antibody staining. Fluorescence imaging was performed in hippocampal CA1 with a LSM780 laser scanning confocal microscope (Zeiss, Zen10 software) using a Plan-Apochromat 40x/NA1.4 oil DIC M27 objective and appropriate excitation laser lines and spectral detection windows.

### Cell culture Imaging

8-bit Z-stacks images (1024 × 1024 pixels) were acquired with a LSM780 confocal microscope (Zeiss) using a Plan-Apochromat 20x/NA0.8 M27 objective. The detector gain in the signal channel(s) was set to cover the full dynamic range but to avoid saturated pixels. Imaging conditions were maintained identical for each experiment.

### Image analysis and representation

For image quantification ten randomly chosen regions of each cell culture dish were imaged and maximum intensity projections were used for quantification. For single cell quantification of FUNCAT, Puromycilation or mRNA signal, single somata were manually outlined based on Map2 staining (to distinguish from glial cells), GFP staining or the relevant marker mRNA (FISH) and areas were saved as ROIs in ImageJ. Then images were split into single channels and the mean intensity value for each ROI was measured in the relevant signal channels. Splitting the neuron population into Cre-negative and Cre-positive neurons within a dish was based on fixed thresholds for FUNCAT signal or GFP signal set by comparison with noANL- and/or Cre-negative culture controls In the protein synthesis inhibitor experiments FUNCAT mean grey values were calculated for each condition from whole images in the signal channel (10 randomly chosen regions per dish). ROI or whole image quantifications were exported from ImageJ, calculations performed in Microsoft Excel and Graphs were plotted in GraphPadPrism. Statistical analysis was performed in GraphPadPrism using ANOVA and correction for multiple comparisons. For quantification of the *mars1* signal in tissue sections, maximum intensity projections of the images from different genotypes were used from 3 sets of mice. Two 100 x 200 µm boxes were placed in each image in a way that the border from stratum pyramidale to stratum radiatum was aligned between the samples and a line profile was calculated in Image J for each box along the longer axis. Profiles of the same genotype were averaged along the Stratum pyramidale/stratum radiatum axis in GraphPad Prism. To estimate the difference between *mars1* signal between Nex-Cre::3xMetRS* and Nex-Cre::1xMetRS* and wild type MetRS we calculated the area under the averaged profile curves for each genotype in GraphPadPrism using the last value as baseline and subtracted the calculated area from the 3xMetRS* and 1xMetRS* without Cre, respectively.

### ANL labeling and administration to living animals

For *in vivo* labeling, 8-to 10-week-old wild type or Cre::floxed-3xMetRS* mice were labeled by a single intraperitoneal injection with 400mM ANL (pH 7.5; 1:10 µL of body weight) and sacrificed at the specified times. Animals were maintained as double homozygous (Nex-Cre::3xMetRS*, DAT-Cre::3xMetRS* and Nex-Cre::1xMetRS*), for 3x and 6x comparison experiments, double heterozygotes were obtained by mating them with wild type animals. At the end of the labeling period, mice were sacrificed with CO_2_ (following local lawfare). For FUNCAT experiments animals were perfused with 4% PFA, treated with sucrose and cut at 30 µm. After click chemistry (performed as described above), fluorescence images were acquired using a microscope (DM 1000, Leica). Sections were photographed with Plan 4X dry objective lens (NA = 0.1) at RT. For BONCAT experiments brain tissue was removed, dissected, and frozen in dry ice.

### BONCAT in tissue

Dissected tissue was homogenized and lysed in PBS supplemented with 1% (w/v) Triton X-100 and 1% (w/v) SDS, along with protease inhibitors (PI, 1:750 dilution of protease inhibitor cocktail 3 w/o EDTA, Calbiochem) and benzonase (Sigma, 1:1,000) at 75 °C for 15 min. Lysates were then cleared by centrifugation and stored at −80 °C until used. BONCAT was performed as previously described (Dieterich et al, 2007). In brief, click chemistry was performed in 120 μL PBS (pH 7.8) supplemented with 0.08% Triton (w/v) SDS, 0.2% (w/v), 90 μg proteins, 300 μM triazole (Sigma, ref. 678937), 50 μM biotin-alkyne tag (Thermo, ref. B10185) or DST (Click Chemistry Tools), and 83 μg/mL CuBr (prepared by dilution of a fresh 10 mg/mL solution in DMSO). The reaction was incubated at 4 °C overnight in the dark. Biotinylated proteins were then separated by electrophoresis and immunoblotted with anti-GFP and anti-biotin antibodies. Tissue samples were lysed as explained before; cleared extracts were alkylated twice with 20 mM iodoacetamide (3 h at RT in the dark) and then cleaned by two passages on PD-SpinTrap G-25 columns (GE Healthcare). Samples were then clicked as explained before. After click chemistry, excess biotin-alkyne was removed by protein precipitation with TCA. Tagged proteins were then affinity-purified with high-capacity Neutravidin agarose beads (Thermo) in PBS supplemented with 1% Triton, 0.15% SDS, and PI (binding buffer) overnight at 4 °C. The beads were then washed extensively at RT, first in binding buffer, then in 0.4% SDS, and finally with PBS + PI and 50 mM ammonium bicarbonate + PI. Bound proteins were eluted by incubation with 5% β-mercaptoethanol, 0.03% SDS, and PI for 30 min at RT. Two consecutive elutions were performed.

### Sample processing for LC-MS/MS

Eluted proteins were prepared for bottom-up proteomics using a suspension trapping protocol as previously reported (Desch et al, 2021). Briefly, samples were mixed with lysis buffer (10% SDS, 100 mM Tris, pH 7.55, with H₃PO₄) in a 1:1 ratio, reduced using 20 mM dithiothreitol for 10 min at RT, and alkylated with 50 mM iodoacetamide for 30 min at RT in the dark. Subsequently, samples were acidified using phosphoric acid to a final concentration of 1.2%. Binding/wash buffer (BW: 90% methanol, 50 mM Tris, pH 7.1 with H₃PO₄) was added in a 1:7 lysate-to-buffer ratio. The protein suspension was loaded onto S-trap filters (size: “micro”; ProtiFi) by centrifugation for 20 s at 4000g. Trapped proteins were washed with 150 μL of BW buffer four times. Trypsin (1 μg; Promega) was added in 60 μL of 40 mM ammonium bicarbonate buffer. Digestion was performed overnight (∼18 hours) at RT in a humidified chamber. Peptides were collected by washing through three consecutive steps by centrifugation at 4000g for 40 s, starting with the digestion buffer and followed by two washes with 0.2% formic acid, in MS-grade water. Peptides were dried *in vacuo* at 45 °C.

### LC-MS/MS analysis

Dried peptides were reconstituted in 95% MS-grade H2O, 5% acetonitrile (ACN), and 0.1% FA. For the time course experiment (3h and 6h ANL labeling), peptides were loaded onto a C18-PepMap 100 trapping column (particle size 3 µm, L = 20 mm, ThermoFisher Scientific) and separated on a C18 analytical column with an integrated emitter (particle size = 1.7 µm, ID = 75 µm, L = 50 cm, CoAnn Technologies) using a nano-HPLC (Dionex U3000 RSLCnano) coupled to a nanoFlex source (2000 V, ThermoFisher Scientific). The temperature of the column oven (SonationAnalytics) was maintained at 55°C. Trapping was carried out for 6 min with a flow rate of 6 μl/min using a loading buffer (100% H2O, 2% ACN with 0.05% trifluoroacetic acid). Peptides were separated by a gradient of water (buffer A: 100% H2O and 0.1% FA) and acetonitrile (buffer B: 80% ACN, 20% H2O, and 0.1% FA) with a constant flow rate of 250 nL/min. In 155 min runs, peptides were eluted by a non-linear gradient with 120 min active gradient time, as selected and reported for the respective MS method by (Muntel *et al*, 2019). Analysis was carried out on a Fusion Lumos mass spectrometer (ThermoFisher Scientific) operated in positive polarity and data-independent acquisition (DIA) mode. In brief, the 40-window DIA method had the following settings: full scan: orbitrap resolution = 120k, AGC target = 125%, mass range = 350-1650 m/z, and maximum injection time = 100 ms. DIA scan: activation type: HCD, HCD collision energy = 27%, orbitrap resolution = 30k, AGC target = 2000%, maximum injection time = dynamic. For the cell type-specific analysis of the olfactory bulb, peptides were analyzed using a nanoElute 1 nano-HPLC coupled to a timsTOF Pro II mass spectrometer via a captive spray ion source (1600 V, Bruker Daltonics). Peptides were loaded directly onto the analytical column (15 cm × 75 µm column with 1.9 μm C18-beads (PepSep); at max. 800 bar), maintained at 60°C and connected to a 20 µm ZDV sprayer (Bruker Daltonics). In 37 min runs, peptides were separated in a linear gradient of water (buffer A: 100% H2O and 0.1% FA) and acetonitrile (buffer B: 100% ACN and 0.1% FA), ramping from 2% to 35% in 30 min with a constant flow rate of 500 nL/min. For data acquisition in DIA mode, the ‘short-gradient’ DIA-PASEF method was employed. In brief, 21 dia-PASEF windows were distributed to a TIMS scan each and designed to cover an m/z range from 475 to 1000 m/z in 25 Da windows, leading to an estimated method cycle time of 0.95 s. The ion mobility range was set from 1.30 to 0.85 Vs/cm². Further details on the method parameters are embedded in the uploaded raw data.

### Data analysis of DIA LC-MS/MS data

DIA raw files were processed with the open-source software DIA-NN (version 1.8.2 beta 27) using a library-free approach. The predicted library was generated using the *in silico* FASTA digest (Trypsin/P) option with the UniProtKB database (Proteome_ID: UP000000589) for Mus musculus. Deep learning-based spectra and RT prediction were enabled. The covered peptide length range was set to 7-35 amino acids, missed cleavages to 2, and precursor charge range to 1-5. N-terminal methionine excision, methionine oxidation, and N-terminal acetylation were set as variable modifications, with cysteine carbamidomethylation as a fixed modification. The maximum number of variable modifications per peptide was limited to 3. According to most of DIA-NN’s default settings, MS1 and MS2 mass accuracies, as well as scan windows, were set to 0; isotopologues and match-between-runs were enabled, while shared spectra were disabled. Protein inference was performed using genes with the heuristic protein inference option enabled. The neural network classifier was set to single-pass mode, and the quantification strategy was selected as ‘QuantUMS (high precision)’. The cross-run normalization was set to ‘RT-dependent’, the library generation to ‘smart profiling’, and the speed and RAM usage to ‘optimal results’. No normalization was applied (additional option “--no-norm”). The DIA-NN report table and the respective FASTA file were imported into the statistical computing software R and analyzed using the MSDAP R package (Koopmans *et al*, 2023). MS-DAP performed protein inference and LFQ quantification at the protein level (min. 1 peptide per protein).

### Data analysis of cell type-specific proteomics

To assess cell type-specific enrichment, ANL-labeled samples were compared to a “no ANL” control. Firstly, a comparison with respect to valid protein quantifications in both conditions (“exclusivity approach”) was performed, and secondly, focusing on relative enrichment over control (“enrichment approach”). For differential abundance analysis, a peptide-centric linear mixed effect model as implemented in the msqRob algorithm in the MS-DAP pipeline was used. Multiple testing correction was performed according to the Benjamini-Hochberg method. For the dopaminergic proteome, proteins were counted as exclusively quantified in the labeled samples (and thus part of the dopaminergic proteome) if 4 of 7 dopaminergic samples and none in the negative control samples had valid protein quantifications. Significantly enriched proteins had to be quantified in at least 4 of 7 labeled and 1 of 5 control samples, have a q-value < 0.01, and show at least 1.5-fold enrichment over control (log2 fold change ∼ 0.58). For the excitatory proteome, proteins were exclusively quantified in the labeled samples (and thus part of the excitatory proteome) if 5 of 6 excitatory samples and none in the negative control samples had valid values. Significantly enriched proteins had to be quantified in at least 5 of 6 labeled and 1 of 2 control samples, have a q-value < 0.01, and show at least 3-fold enrichment over control (log2 fold change ∼ 1.58). To assess changes in protein intensity after prolonged labeling duration, the DIA-NN precursor report was filtered for proteins matching the intersection of the previously identified excitatory proteome of the two time points, before performing the linear mixed effect model in MS-DAP (q-value < 0.01, 1.5-fold enrichment).To compare the dopaminergic and excitatory neuronal proteomes to previously published studies, a gene-level approach was used to integrate the different datasets. Before merging, any information on protein isoforms (e.g., protein half-life) was aggregated to gene IDs by their mean value. Gene ontology and PANTHER pathway overrepresentation analyses were performed using ShinyGO v0.80 (Ge *et al*, 2020) with a custom background list of the overall olfactory bulb proteome derived from Sharma et al. (Sharma *et al*., 2015), applying an FDR cutoff of 0.05 and plotting in order of lowest FDR (top terms). All visualizations were performed using base R or the ggplot2 package.

## Data availability

All mass spectrometry proteomics raw data and associated search engine results have been deposited to the ProteomeXchange Consortium via the PRIDE partner repository (Perez-Riverol *et al*, 2019) with the dataset identifier PXD057536. Analysis scripts are available upon request.

## Acknowledgements

We thank Mariam Rahmatian, Ina Bartnik, Nicole Fürst and Elena Ciirdaeva for excellent technical assistance and Emily Northrup, Nataliya Golovyashkina, Nina Vogt and Silke Zeissler and the animal facility of MPI for Brain Research for their excellent support. We thank Bachelor student Ainhoa Gallego Alguacil and the intern Anna Pfeifer for their help with some of the experiments and Miguel Diaz-Hernandez for his general support. BAC is funded by the Spanish Ministry of Science and Innovation (Ramón y Cajal-RYC2018-024435-I), by the Autonomous Community of Madrid (Atracción de Talento-2019T1/BMD-14057), and MICINN (PID2020-113270RA-I00) grants. RAP is funded by Autonomous Community of Madrid (Atracción de Talento-2019T1/BMD-14057). EMS is funded by the Max Planck Society, and the European Commission (Advanced ERC DiverseSynapse, 101054512).

## Author contributions

BAC, EMS and StD designed the experiments. BAC, StD, KD, RAP, BNA, RSS, COS acquired, analyzed and interpreted the data. BAC, StD, KD, JDL and EMS analyzed and interpreted the data. BAC, StD, KD and EMS wrote the manuscript. All authors contributed to the writing and revision of the article.

---

**Supplementary Figure 1.**
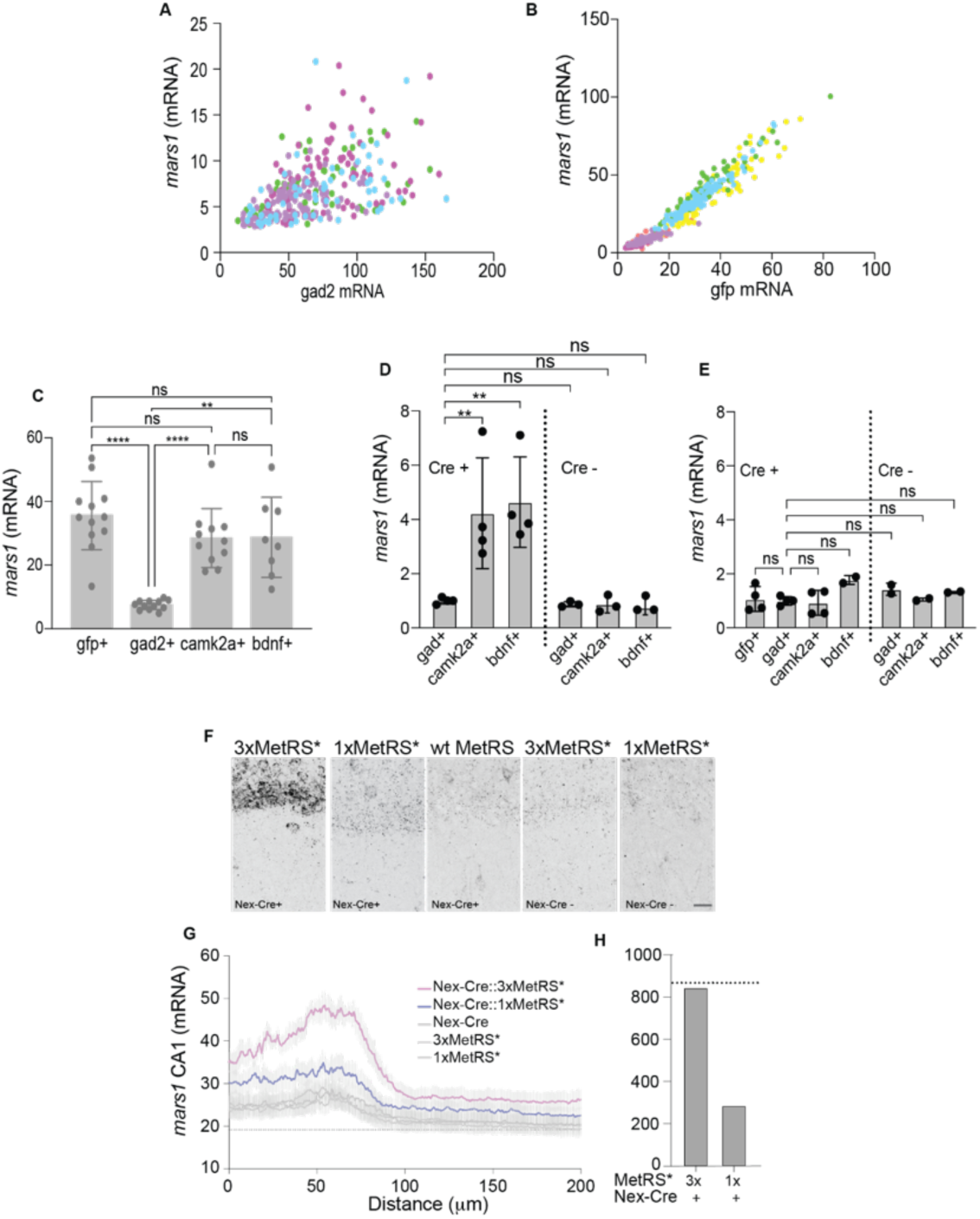
FISH for MetRS mRNA (*mars1)* in cultured neurons and hippocampal slices. **(A),** FISH signals for *mars1* and *gad2* (mRNA) are distributed in the same intensity space in inhibitory cells (*gad2+*) from Nex-Cre::3xMetRS* (blue, green dots) and Nex-Cre::1xMetRS* (pink, purple dots) cultures. (**B),** Both, mRNA FISH signals for *gfp* and *mars1* are substantially higher in *gfp+* cells from Nex-Cre::3xMetRS* (blue, green, yellows) than from Nex-Cre::1xMetRS* (pink, purple, reds) cultures. (**C),** Quantification of *mars1* in different cell types identified by FISH signal (*gfp+, gad2+*, *camk2a+* or *bdnf+*,) within cultured neurons from Nex-Cre::3xMetRS* mice. Excitatory cells (*camk2a+, bdnf+*) have comparable *mars1* signal to *gfp+* neurons while inhibitory cells (*gad2+*) have lower *mars1* signal, **** p<0.0001, ** p<0.0091. **D, E** Quantification of *mars1* in different cell types identified by FISH signal (*gad2+*, *camk2a+* or *bdnf+*) in primary cultured neurons from 3xMetRS*, ** p<0.0080 (gad+/camk2a+), ** p<0.0030 (gad+/bdnf+), (D) or 1xMetRS* (E) mice expressing Nex-Cre (left part) or not (right part) show significant overexpression of *mars1* only in Nex-Cre::3xMetRS* excitatory cells. The signal is normalized on *mars1* in *gad+* neurons from Nex-Cre-positive cultures. **F**, Confocal images of FISH detecting MetRS mRNA in the CA1 layer of the hippocampus in brain slices from 1xMetRS* and 3xMetRS* mice. Scale bar 25 µm. **G**, Quantification of MetRS mRNA as a function of the distance to the CA1 pyramidal layer shown in F. **H**, Quantification of the area under the curve for MetRS* mRNA expression shown in (F).

**Supplementary Figure 2.**
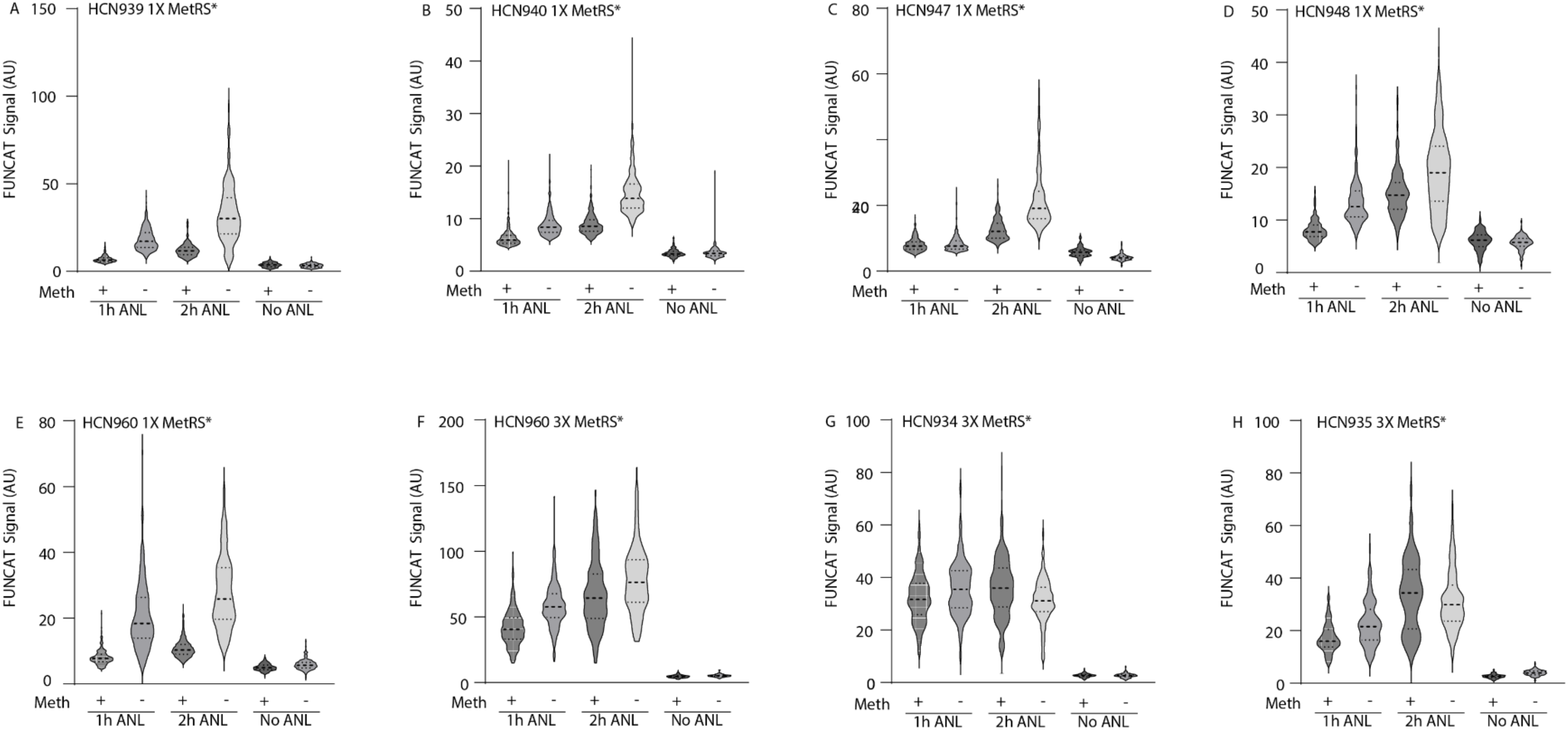
Methionine competition with ANL incorporation to MetRS*. Violin plots representing the FUNCAT quantification of each neuron from the primary neuronal cultures shown in Fig. 3, from Nex-Cre::1xMetRS* and Nex-Cre::3xMetRS* mice, labeled with ANL in culture media with (+) or without (-) methionine for 1 hour and 2 hours. In each graph (A-H), a single experiment is represented.

**Supplementary Figure 3.**
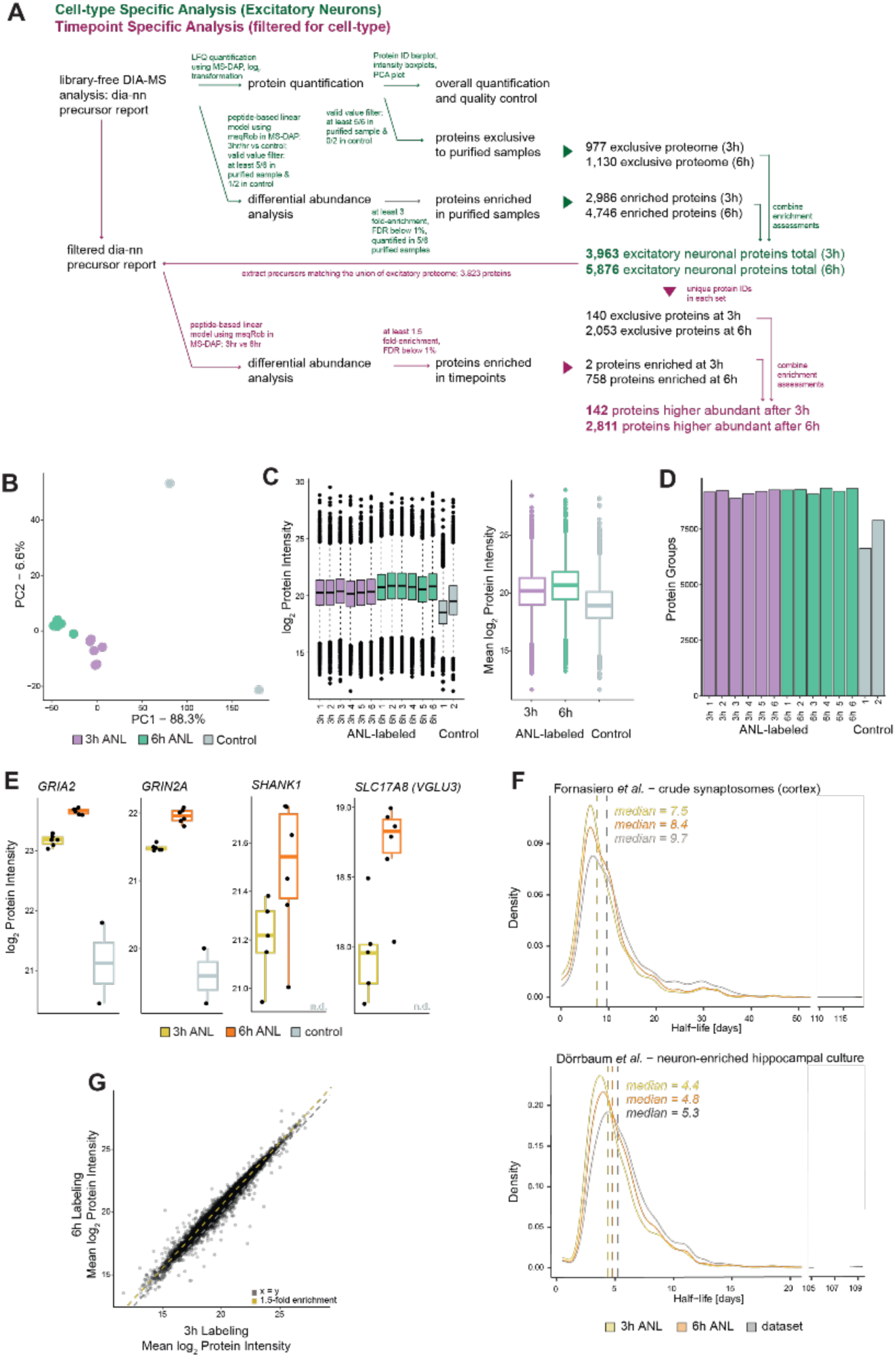
Proteomic analysis of excitatory neuronal proteins from mouse cortex using short labeling *in vivo*. (**A**) Detailed bioinformatic analysis pipeline for the characterization of the excitatory neuronal proteome from the cortex after 3 hours and 6 hours of labeling with ANL *in vivo*. All relevant steps of the two-staged analysis—cell type specificity (Hrvatin *et al*.) and temporal specificity (maroon)—are indicated. (**B**) Principal component analysis (PCA) of overall protein group quantifications (log2) showing clustering of biological replicates and separation of conditions, indicating similarity between the proteins after 3 hours and 6 hours of labeling (>88% variance explained by component 1). (**C**) Log2-scaled protein intensities of the identified proteins for each biological replicate (left) and log2 mean intensities for each time point and control samples (right). (**D**) Box plots of overall protein group intensities (log2) for each condition, separated by biological replicates (left) or merged by mean values (right). Boxes indicate the median, first, and third quartiles, and whiskers extend to 1.5 × IQR. (**E**) Box plots display selected excitatory candidate proteins showing either significant enrichment in the labeled over the unlabeled control samples (GRIA2, GRIN2A) or exclusive detection in the labeled conditions (VGLU3, SHANK1). Boxes indicate the median, first, and third quartiles, and whiskers extend to 1.5 × IQR. These proteins are examples of differential abundances at 3 hours or 6 hours (as shown in Fig. 6H). (**F**) The density plot shows protein half-life distribution in mouse cortical synaptosomes (left) or neuron-enriched hippocampal cultures (right), (Dorrbaum *et al*., 2018). Excitatory neuronal proteins identified at each time point are matched to previously published databases at the level of gene symbols and are highlighted by color; overall half-life data is depicted in gray. Vertical lines in the histogram highlight lower median half-lives of the matched excitatory proteins after 3 hours than after 6 hours, both having shorter half-lives compared to the full proteome. (**G**) Scatter plot of mean log2 of protein intensities of all identified proteins at 3 hours and 6 hours, showing a shift towards higher intensities at 6 hours. See also panel C (right).

**Supplementary Figure 4.**
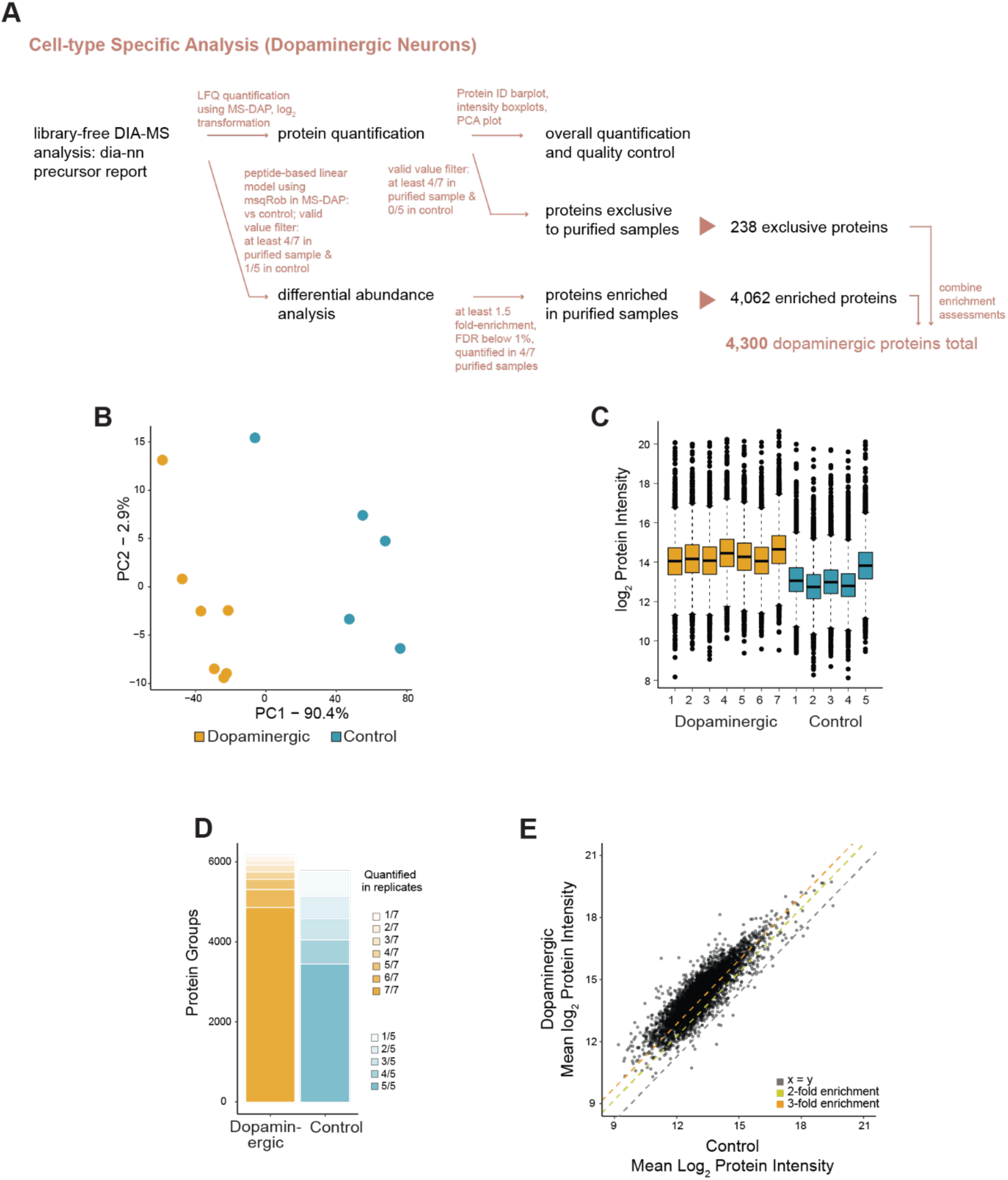
Proteomic analysis of the dopaminergic neuronal proteome of the olfactory bulb using *in vivo* labeling. (**A**) Detailed bioinformatic analysis pipeline for the characterization of the OB-DA proteome after two weeks of labeling by daily IP injections of ANL. All relevant steps of the cell type-specific analysis are indicated. (**B**) Principal component analysis of overall protein group quantifications (log2) showing separation of the labeled vs. unlabeled samples (<90.4% variance explained by component 1) and clustering of biological replicates. (**C**) Boxplots of overall protein group intensities (log2) for each condition, separated by biological replicates. Boxes specify the median, first and third quantile, and whiskers extend to 1.5 × IQR. (**D**) Bar plot of overall protein quantifications for the labeled dopaminergic and unlabeled control samples. Saturation of the stacked bars indicates the number of valid observations for a protein group in all biological replicates. (**E**) Scatterplot of mean log2 protein group intensities of all identified proteins in the unlabeled negative control vs. labeled samples, showing a shift toward higher intensities for the labeled dopaminergic proteins. See also panel C.

